# Enabling high-accuracy long-read amplicon sequences using unique molecular identifiers with Nanopore or PacBio sequencing

**DOI:** 10.1101/645903

**Authors:** Søren M. Karst, Ryan M. Ziels, Rasmus H. Kirkegaard, Emil A. Sørensen, Daniel McDonald, Qiyun Zhu, Rob Knight, Mads Albertsen

**Author notes:** These authors contributed equally to this work (Søren M. Karst, Ryan M. Ziels). Correspondence: Mads Albertsen.

## Abstract

High-throughput amplicon sequencing of large genomic regions remains challenging for short-read technologies. Here, we report a high-throughput amplicon sequencing approach combining unique molecular identifiers (UMIs) with Oxford Nanopore Technologies or Pacific Biosciences CCS sequencing, yielding high accuracy single-molecule consensus sequences of large genomic regions. Our approach generates amplicon and genomic sequences of >10,000 bp in length with a mean error-rate of 0.0049-0.0006% and chimera rate <0.022%.

## Main

High-throughput amplicon sequencing is a ubiquitous method for studying genetic populations with low-abundance variants or high heterogeneity, including cancer driver genes^1–3^, virus populations^4–6^ and microbial communities^7^. Short-read Illumina sequencing has dominated amplicon-related research due to its unprecedented throughput and low native error-rate of ∼0.1%, but its maximum amplicon size of ∼500 bp^8^ limits important long-range information and assay resolution^9^. Unique molecular identifiers (UMIs) have been applied to enable sequencing of longer amplicons with short-reads via assembly of synthetic long reads^10^. Each template molecule in a sample is tagged with a UMI sequence consisting of 10-20 random bases, which can subsequently be used to sort and analyse reads based on their original template molecule. UMIs can enable sequencing of synthetic long-reads up to ∼11,000 bp, but this approach cannot resolve amplicons with repeats longer than the short-read length^11^, which limits its application. The high native error rates of Oxford Nanopore Technologies (ONT) (5-25%^12^) and Pacific Biosciences (PacBio) (13%^13^) have, until now, made it difficult to confidently identify true UMI tag sequences necessary to accurately assign raw reads to their template molecules. Furthermore, the combination of UMIs with long-read sequencing is relatively unexplored, and only recently has it been applied with PacBio CCS^14–16^, but without using dual UMIs for chimera filtering^17^ and profiling the error of the generated consensus sequences.

Here, we present a simple workflow that combines UMIs with sequencing of long amplicons on the ONT and PacBio platforms to produce highly accurate single-molecule consensus sequences with a low chimera rate. To improve recognition of UMI-tagged error-prone reads, we designed UMIs to contain recognizable internal patterns (**Figure 1C** and **Table S1**) that avoid error-prone homopolymer stretches^18^, which combined with filtering based on UMI length and pattern allows for robust determination of true UMI sequences in raw error-prone ONT and PacBio data.

**Figure 1:**
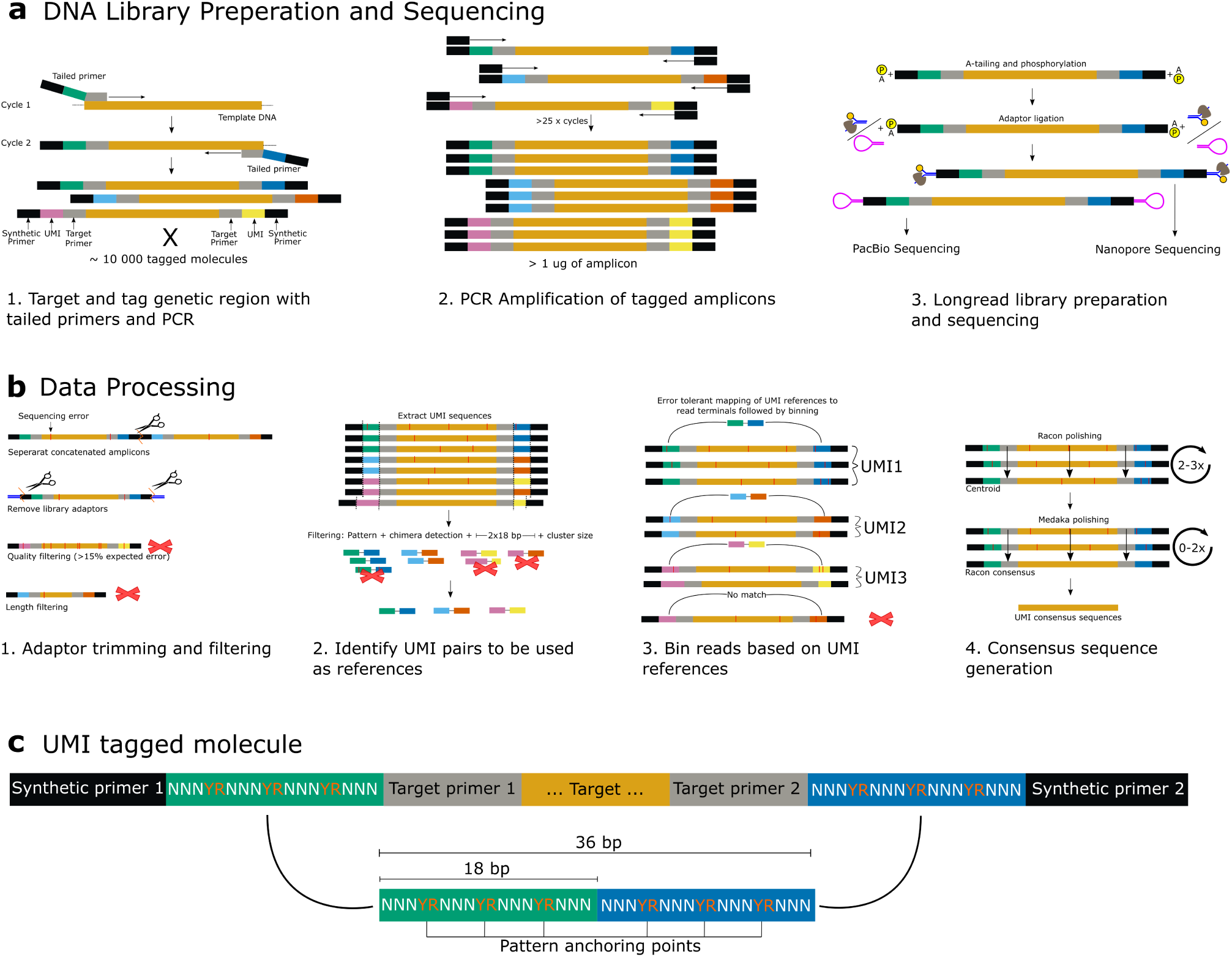
Overview of laboratory (a) and bioinformatics workflow (b). (c) A schematic overview of the dual UMI-tagged molecule. The two UMIs are detected and processed together in the bioinformatics pipeline.

The DNA template is initially diluted to the target number of output sequences, which is estimated based on the desired single-molecule coverage and expected sequencing yield. The genetic region of interest is then targeted using 2 cycles of PCR with a customized set of tailed primers (Table S1), which includes a target-specific primer, a UMI sequence and a synthetic priming site used for downstream amplification (Figure 1A, step 1). For PCR-free approaches, such as whole genome or metagenomic DNA sequencing, the adapters containing UMIs can be ligated to the template DNA molecules. The product from the initial UMI-tagging step is a double-stranded DNA amplicon copy of the genetic target, containing the UMIs and synthetic primer sites on both ends. The UMI-tagged molecule is subsequently amplified by PCR targeting the synthetic primer sites (Figure 1A, step 2), and prepared for long-read sequencing with ONT or PacBio CCS (Figure 1A, step 3). After sequencing, reads are binned based on both terminal UMIs (Figure 1B, steps 1 and 2). UMI sequences that have a high probability of being correct are detected based on the presence of a designated pattern, as well as an expected UMI length of 18 bp. Chimeric sequences are *de novo* filtered by UMI-pairs in which either terminal UMI is observed in a more abundant UMI-pair^17^ (Figure 1B, step 2). The filtered, high-quality UMI-pair sequences are used as a reference for binning the raw dataset according to UMIs (Figure 1B, step 3). The consensus sequence for each UMI bin is then generated by multiple rounds of polishing using the binned raw reads (Figure 1B, step 4).

To assess the effectiveness of our UMI-tagging approach for error-correcting long reads, we sequenced full-length ribosomal RNA (rRNA) operons (∼4400 bp) in a mock microbial community from ZymoBIOMICS containing eight bacterial species (**Table S13**). To compare sequencing approaches, we calculated the necessary read coverage to obtain a mean error rate < 0.01%/Q40 (**Table S2**), and data below that read coverage cut-off was removed from the analysis. Afterwards, chimeric sequences and sequences from reagent contamination were identified (**Figure S8**), manually curated, and removed from the dataset to enable calculations of true error-rates. On a single ONT MinION R10 flowcell, a total of 23,365 amplicon UMI consensus sequences (ONT UMI) were generated with read coverages ≥ 25x (Q40 cutoff), an average length of 4,381 bp, a mean residual error rate of 0.0049%, and a chimera rate of 0.017% (**Figure 2a, b**). Sequencing the same library of UMI-tagged long amplicons with a PacBio Sequel II 8M flowcell in CCS mode resulted in 39,678 UMI consensus sequences (PB UMI) with read coverages ≥ 3x (Q40 cutoff), an average length of 4,376bp, a mean residual error rate of 0.0006%, and a chimera rate of 0.022% (**Figure 2a, b**). For comparison, raw PacBio CCS reads without UMI clustering generated 135,823 CCS reads (PB CCS) with read coverages ≥ 40x (Q40 cutoff), which had a mean error rate of 0.0080% and a chimera rate of 1.9%. The 1.9% chimeras observed in the raw PB CCS data are most likely introduced during the PCR amplification of the UMI library, and are therefore present in the amplicon library before sequencing. The exact same amplicon library was used to produce the ONT UMI and PB UMI data. Thus, using a rigorous UMI-based filtering approach almost eliminates PCR chimeras, which otherwise can make up over 20% of the amplicons depending on the PCR conditions^19^. The ligation-based ONT UMI approach was tested on genomic DNA fragments of up to 10,000 bp from *Escherichia coli*, and the results were consistent with the rRNA operon results (**Figure S3**), demonstrating the flexibility of this method for improving the accuracy of long-read sequencing.

**Figure 2:**
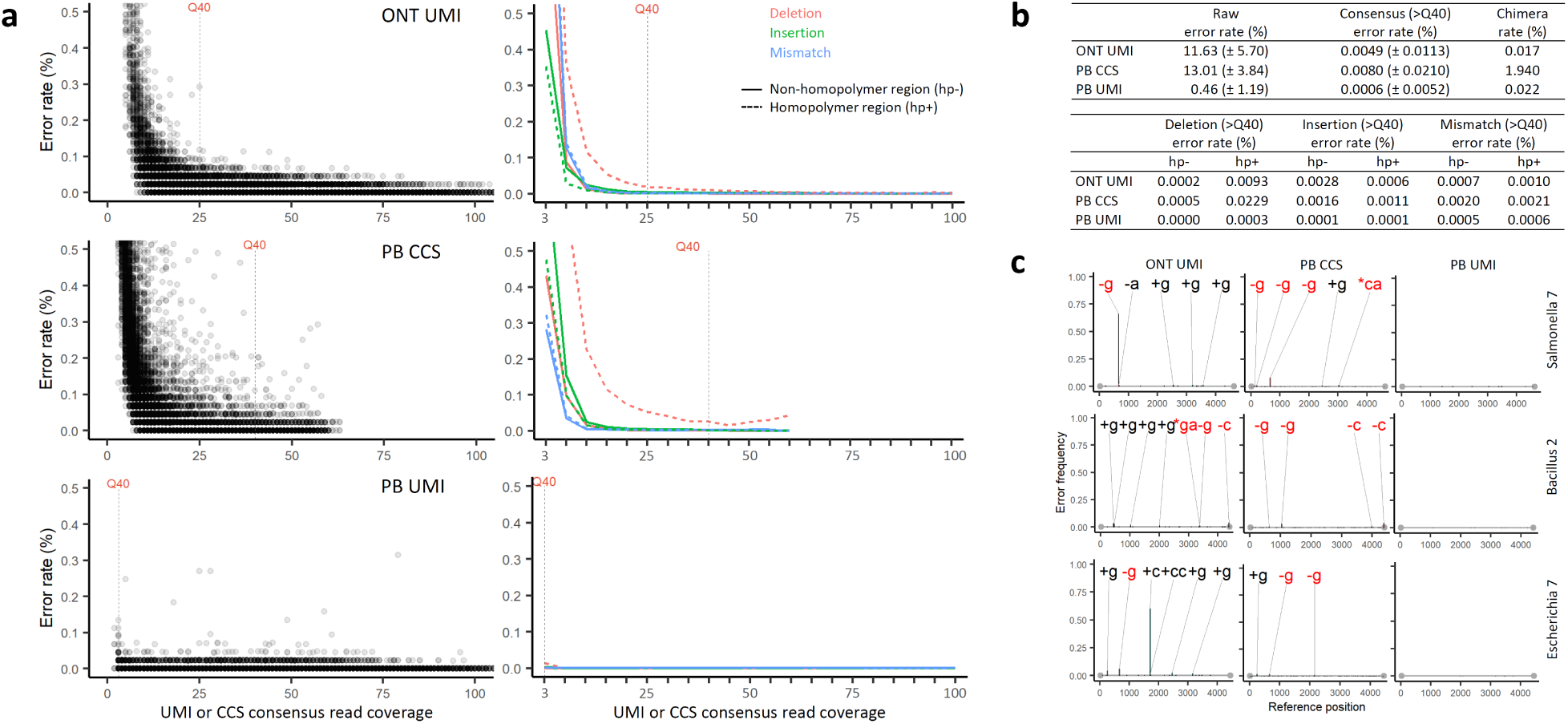
a) Left column: Error rate as a function of the number of reads in each UMI bin for the three data types. Right column: Error rate as a function of the number of reads in each UMI bin split by error type and whether the error fell inside (hp+) or outside (hp-) a homopolymer region. b) The top table shows the mean error rate (+/- standard deviation) of raw reads and consensus sequences (CCS/UMI) with a Q40 minimum and the observed chimera rate. The bottom table summarises the mean error rates for all error and homopolymer types for data with a Q40 minimum. c) Frequency of specific errors are plotted as a function of operon position (bp) for Salmonella operon 7, Bacillus operon 2 and Escherichia operon 7 for ONT UMI, PB CCS and PB UMI respectively. The error frequency is normalized as fractions of sequences containing the error in that position. Errors with ≥ 0.01 frequency have been annotated with error type. +[actg] means insertion -[actg] means deletion and *[actg][actg] means mismatch. Annotated errors in black are in non-homopolymer regions and errors in red are in homopolymer regions.

The residual errors were markedly different in non-homopolymer regions compared to homopolymer regions for both the ONT UMI and PB CCS data, while the PB UMI data was extremely low in both cases (**Figure 2a, c**). The error rate for all error types (deletions, insertions, and mismatches), except homopolymer deletions, stabilized for both ONT UMI and PB CCS above a coverage of 20x (**Figure 2a, Table S2**). The high deletion error rate was primarily due to deletions in long (>4 bp) C and G homopolymers for both data types (**Figure S4-5**, **Table S3**), and reaffirmed that homopolymer-derived errors are a remaining obstacle for lower error rates. For the ONT UMI data, G-insertions in non-homopolymer regions made up the majority of remaining errors. Both the non-homopolymer insertions and the homopolymer deletions were to some degree systematic, with some errors in specific positions being present in >50% of the sequences (**Figure 2c** and **Figure S4**). For PB CCS data, the homopolymer deletion errors were not as systematic, but still a major error contributor (**Figure 2b, Figure S5**). Random mismatch error is the other major source of error for PB CCS data, which probably originate from PCR errors. For the PB UMI data, there are very few errors left (1109 errors in 39678 sequences of ∼4400 bp), and thus not enough data to elucidate potential error trends (**Figure S6, Table S3**).

Characterizing whether the residual error is random or systematic is important for the ability to accurately call variants from single molecule consensus sequences. We naively generated variants from the three data types (data >Q40 threshold but without removing chimeras) by clustering consensus sequences, phasing single nucleotide variants (SNVs) within clusters and calling variants if present at ≥3x coverage. The 43 references in the Zymo Mock were observed for all datatypes, (**Figure S7**), with no errors except for 2 variants from the ONT UMI data, which each had 1 error in homopolymer regions (**Figure S4**). For ONT UMI, PB CCS and PB UMI data, 1.00%/6.99%/0.18% of the consensus data were respectively assigned to variants with systematic errors, and 0%/0.46%/0% were assigned to chimeric variants (**Table S4**).

CCS-like strategies have been attempted as an alternative to UMIs to reduce the error-rate of amplicon sequencing on the ONT platform, but these methods suffer from insufficient molecule coverage to effectively reduce mean error rates below 2%^20^ (**Table S10**). In principle, lower error rates can also be achieved with denoising strategies^20, 21^, but at the cost of potentially missing true low-abundance variants^22^, which are critical for some applications (e.g. pathogen detection and drug resistance). Furthermore, state-of-the-art clustering algorithms depend on the abundance of unique sequences to model errors^23, 24^, which is not suitable for datasets where population micro-heterogeneity is high and evenness low.

Amplification and sequencing of rRNA genes has become an important method for studying the diversity and taxonomic composition of human- and environment-associated microbial communities. Here, we applied the PB UMI method to generate 253,089 high-quality, full-length bacterial rRNA operon sequences from 70 human fecal samples collected by the American Gut Project^25^. We assessed strain-level taxonomic resolution by annotating the full length 5S, 16S and 23S within the operons, and searched for these genes against gene-specific databases from the “Web of Life” ^26^. Using only the full-length 16S rRNA gene, 11.3% of the sequences could be matched at the strain level to the database, and 38.4% assigned at the species level. By using both the 16S and 23S rRNA genes within an operon, we could assign 22% at the strain level and 72.2% at the species level, representing a significant increase in assignment over using the full length 16S rRNA alone (Chi-squared statistic=124,086, p-value<2.23e-308; **Figure 3**). These results are inline with a recent study of the taxonomic resolution of the rRNA operon^27^. This UMI approach should also enable direct quantification of molecules in a sample^28^, which would be ideal for precise relative abundance estimates. However, the mock community used here, and probably many others, contains a biased fragment size and growth-dependent coverage, preventing proper quantification (**Table S5 and Figure S9-13**).

**Figure 3:**
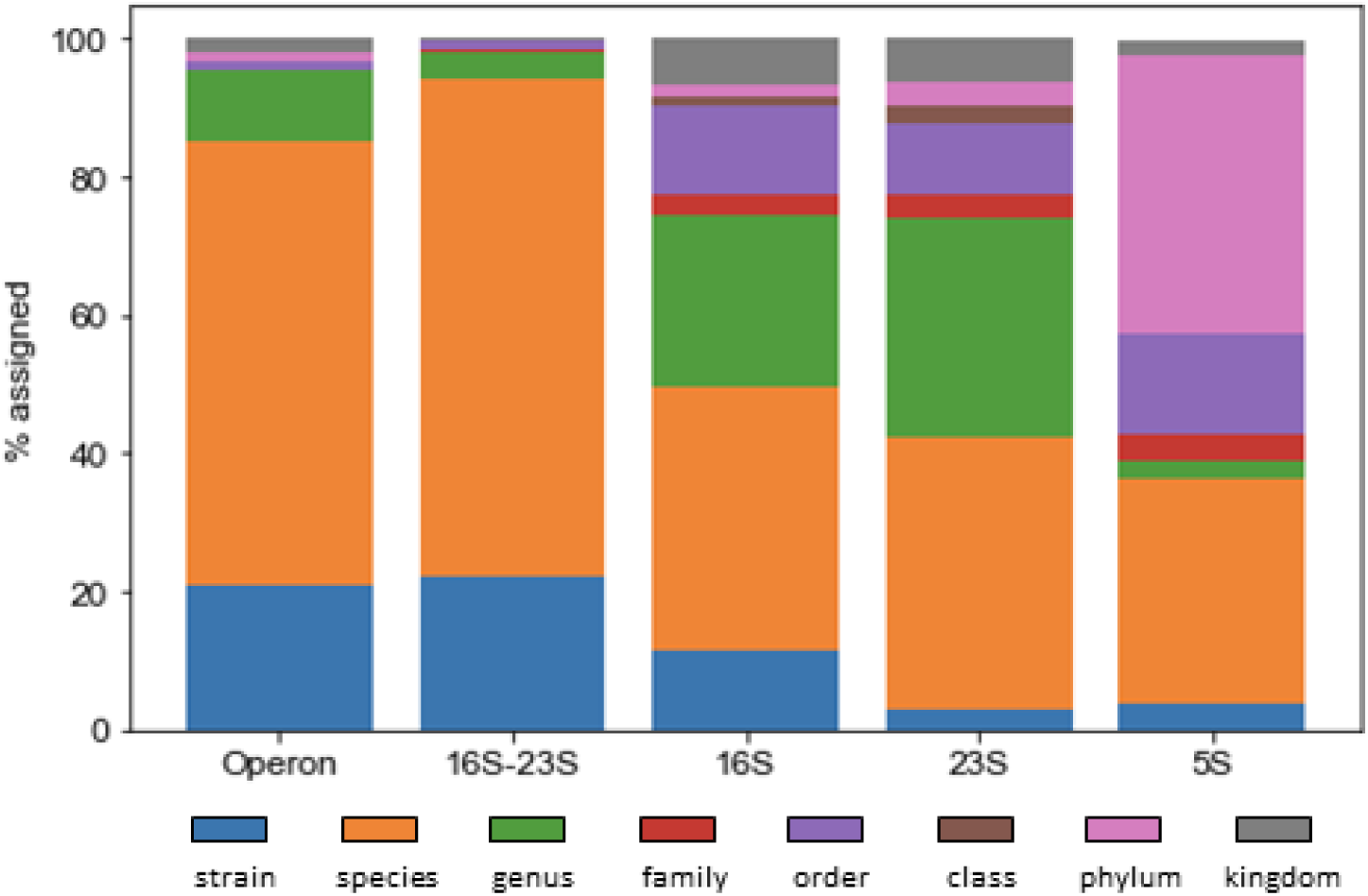
BLAST-based consensus taxonomic assignment against the Web of Life 86k reference database for whole rRNA operons, using the combination of 16S and 23S rRNAs, and the individual rRNA genes. In the dataset, 253,089 operons were available and used for assignment. Of these, n=253,087 had an annotatable 23S rRNA gene, n=253,088 had annotatable 16S rRNA gene, and n=50,560 had annotatable 5S rRNA gene. All raw and annotatable elements were used in this summary.

The choice of library strategy and sequencing platform for high-accuracy amplicon sequencing depends on the application. An overview of time, cost and yield comparisons was compiled for the three different approaches (**Table S6-8**), and the current projected price per sequence for >Q40 data from rRNA operons is: 0.015 USD (ONT UMI), 0.012 USD (PB CCS), 0.007 USD (PB UMI), which should be similar for other genetic targets. For rapid testing and iterative development, the ONT UMI approach is attractive due to its low cost and portability. PB CCS sequencing also performs well for high-accuracy amplicon sequencing, but the presence of low abundant chimeric variants is problematic, especially if they propagate into reference databases^29^. For sensitive applications, such as detecting low-abundance variants or generating reference sequences for key databases, the PB UMI approach appears to be the best suited at this time.

## Acknowledgements

The study was funded by research grants from VILLUM FONDEN (15510) and the Poul Due Jensen Foundation (Microflora Danica). RMZ was funded by grants from the Natural Sciences and Engineering Research Council of Canada (Discovery Grant), and Genome British Columbia (SIP011).

## Contributions

SMK and RMZ conceived the method and developed the bioinformatics pipeline. SMK performed wet lab method development and experiments. EASO performed Nanopore UMI sequencing of E. coli. RHK assembled reference genomes. SMK, RMZ and MA performed data analysis on method performance. DM, QZ, RK analysed American Gut Project samples. SMK, RMZ, MA wrote the first draft of the manuscript. All authors contributed to the content and revision of the manuscript.

## Methods

### rRNA operon UMI sequencing of mock microbial community

#### Source of DNA

The ZymoBIOMICS Microbial Community DNA Standard (D6306, lot no. ZRC190811) was obtained from Zymo Research (Irvine, California). The mock community DNA contained genomic material from 10 species (8 bacteria and 2 yeasts): *Bacillus subtilis*, *Cryptococcus neoformans*, *Enterococcus faecalis*, *Escherichia coli*, *Lactobacillus fermentum*, *Listeria monocytogenes*, *Pseudomonas aeruginosa*, *Saccharomyces cerevisiae*, *Salmonella enterica*, *Staphylococcus aureus*. Note, 2 of the yeast species were not targeted by PCR amplification of rRNA operons based on the primers used (see *DNA Sequence Library Preparation*). The concentration of DNA in the mock community sample was measured on a Qubit 3.0 fluorometer and Qubit dsDNA HS assay kit (Thermo Fisher Scientific), and the quality of the DNA was measured by gel electrophoresis on an Agilent 2200 Tapestation using Genomic Screentapes (Agilent Technologies).

#### DNA Sequence Library Preparation

##### Online protocol

An interactive step-by-step protocol is available at protocols.io: (https://www.protocols.io/private/F5C5FE21305911EAAC0B0242AC110003).

##### Tagging target gene with UMIs

PCR was used to target the bacterial 16S-23S rRNA operon and simultaneously tag each template molecule with terminal unique molecular identifiers (UMIs).

The following tailed primers lu_16S_8F_v7 and lu_23S_2490R_v7 were used for the PCR (see **Supplementary Table 1**). The first section of both tailed primers is a synthetic priming site used for downstream amplification. The second section is the ‘patterned’ UMI consisting of a total of 12 random nucleotides (N) and 6 degenerate nucleotides (Y or R) which results in a total of 1.2×10^18^ possible UMI combinations when UMIs from both terminal ends of a template molecule are concatenated (4^12*2^ × 2^6*2^ = 1.2×10^18^, see **Figure 1**). The last section of the tailed primers consists of the rRNA operon-specific primer for 27f^1^ or 2490r^2^, respectively.

The PCR reaction contained 5 ng of ZymoBIOMICS Microbial Community DNA Standard, 1 U Platinum SuperFi DNA Polymerase High Fidelity (Thermo Fisher Scientific, USA), and a final concentration of 1X SuperFi buffer, 0.2 mM of each dNTP, 500 nM of each lu_16S_8F_v7/ lu_23S_2490R_v7 primers in 50 µL. The PCR program consisted of initial denaturation (3 minutes at 95°C) and 2 cycles of denaturation (30 seconds at 95°C), annealing (30 seconds at 55°C) and extension (6 minutes at 72°C). The PCR product was purified using a custom bead purification protocol “SPRI size selection protocol for >1.5-2 kb DNA fragments” (Oxford Nanopore, England) based on: dx.doi.org/10.17504/protocols.io.idmca46. CleanPCR (CleanNA, Netherlands) bead solution was used for preparing the custom buffer. The purification was performed according to the custom protocol with the exception of an EtOH concentration of 80% and 0.9x bead solution/sample ratio.

##### Amplification of UMI-tagged amplicons

A second PCR was used to amplify the UMI-tagged template molecules. All of the UMI-tagged template molecules were added to the reaction containing 2 U Platinum SuperFi DNA Polymerase High Fidelity (Thermo Fisher Scientific, USA), and a final concentration of 1X SuperFi buffer, 0.2 mM of each dNTP, 500 nM of each lu_pcr_fw_v7 and lu_pcr_rv_v7 primers (see **Table S1**) in 100 µL. The PCR program consisted of initial denaturation (3 minutes at 95°C) and then 25 cycles of denaturation (15 seconds at 95°C), annealing (30 seconds at 60°C) and extension (6 minutes at 72°C) followed by final extension (5 minutes at 72°C). The PCR product was purified using a custom bead purification protocol “SPRI size selection protocol for >1.5-2 kb DNA fragments” (Oxford Nanopore, England) based on: dx.doi.org/10.17504/protocols.io.idmca. CleanPCR (CleanNA, Netherlands) bead solution was used for preparing the custom buffer. The purification was performed according to the custom protocol with the exception of an EtOH concentration of 80% and 0.9x bead solution/sample ratio. The concentration and quality of the PCR amplicons was measured as described before.

To obtain sufficient PCR product for sequencing, a third PCR was performed using the amplicons generated from the second PCR and using the same procedure as before, but with 3 × 100 µl reactions and 6 cycles of amplification. A large reaction volume was used to minimize the risk of overamplification. The final amount of amplicon DNA generated was 3.5 µg in 55 µL.

#### Oxford Nanopore sequencing of mock rRNA operon amplicons

1200 ng of the purified amplicon DNA from the third PCR was used as template for library preparation using the protocol “1D amplicon/cDNA by ligation (version ACDE_9064_v109_revA_23May2018, SQK-LSK109)” (Oxford Nanopore, England). A MinION R10 flowcell (FLO-MIN110) was used for sequencing on a MinION and MinION software v19.10.1 (Oxford Nanopore, England). Basecalling was performed with Guppy v3.4.4 (Oxford Nanopore, England) in GPU mode with following modifications to the standard settings *--trim_strategy ‘none’ --device cuda:0 --chunk_size 1500 --chunks_per_runner 1024 --config dna_r10_450bps_hac.cfg model*.

#### PacBio Sequel II sequencing of mock rRNA operon amplicons

2000 ng of the purified amplicon DNA from the third PCR was sent for PacBio library preparation and sequencing at the DNA Sequencing Center at Brigham Young University (https://dnasc.byu.edu/). The amplicons were incubated with T4 polynucleotide kinase (New England Biolabs, USA), following the manufacturer’s instructions. The sequencing library was prepared using SMRTbell Express Template Preparation Kit 1.0 following the standard protocol. Sequencing was performed on a Sequel II using Sequel II Sequencing Kit 1.0, Sequel II Binding and Int Ctrl Kit 1.0 and Sequel II SMRT Cell 8M following standard protocol with 1 hour pre-extension and 30 hour collection time (Pacific Biosciences, USA). Circular consensus (CCS) reads were generated using CCS version 3.4.1 (https://github.com/PacificBiosciences/ccs) using standard settings.

#### Data generation workflow

##### Online scripts

Source code and analysis scripts are freely available at https://github.com/SorenKarst/longread-UMI-pipeline

##### Trimming and filtering of raw data (Nanopore data only)

Raw fastq sequence data was trimmed of sequencing adapters using porechop with the commands: --*min_split_read_size 3500 --adaptor_threshold 80 --min_trim_size 20 --extra_end_trim 0 --extra_middle_trim_good_side 0 --extra_middle_trim_bad_side 0--middle_threshold 80 --check_reads 5000* (v0.2.4 https://github.com/rrwick/Porechop). Additionally, the *adaptors.py* file in porechop was modified to include possible end-to-end ligation combinations of the customized primers. The customized settings and modifications to the *adaptors.py* file were necessary to correctly split amplicons concatenated in the ligation step of the library preparation, which can make up a substantial amount of the data.

The adaptor trimmed data was filtered using filtlong with the settings *--min_length 3500--min_mean_q 70* (v0.2.0 https://github.com/rrwick/Filtlong) and cutadapt^3^ (v2.7) with *-m 3500–M 6000*. The output from these pre-processing steps was trimmed and filtered raw read data.

##### Extraction of UMI reference sequences

To efficiently bin reads according to the UMIs on their terminal ends, it was critical to extract and validate true UMI sequences that could be used as references for subsequent mapping steps. UMI sequences of the correct length (18 bp) were extracted from the reads by locating the flanking sequences within the custom primers. To do so, the first 200 bp from each terminal end of all reads were extracted, and saved into individual files. UMI sequences were extracted from each terminal end file with *cutadapt -e 0.2 -O 11 -m 18 -M 18 --discard-untrimmed*

*-g CAAGCAGAAGACGGCATACGAGAT…AGRGTTYGATYMTGGCTCAG*

*-g AATGATACGGCGACCACCGAGATC…CGACATCGAGGTGCCAAAC*

*-G GTTTGGCACCTCGATGTCG…GATCTCGGTGGTCGCCGTATCATT*

*-G CTGAGCCAKRATCRAACYCT…ATCTCGTATGCCGTCTTCTGCTTG* in paired-end input mode. This step insured that only reads with UMIs of the correct length (18 bp) in both ends were extracted. UMI pairs were then concatenated and filtered with *grep -B1 -E “NNNYRNNNYRNNNYRNNNNNNYRNNNYRNNNYRNNN”* to remove UMI pairs that did not follow the expected pattern. Filtered UMI pairs were clustered using usearch^4^ (v11.0.667) with the commands *usearch -fastx_uniques -minuniquesize 1 -strand both* and *usearch -cluster_fast-id 0.90 -centroids -sizein -sizeout -strand both -minsize 1*. To estimate the coverage of the clustered UMI pairs, UMI sequences were re-extracted using only a single primer with *cutadapt -e 0.2 -O 11 -m 18 -L 18 --discard-untrimmed*

*-g CAAGCAGAAGACGGCATACGAGAT -g AATGATACGGCGACCACCGAGATC*

*-G GATCTCGGTGGTCGCCGTATCATT -G ATCTCGTATGCCGTCTTCTGCTTG* in paired-end input mode. The re-extracted UMI pairs were concatenated and mapped to the clustered UMI pairs using bwa^5^(v0.7.17-r1198-dirty) with the commands: *bwa index, bwa aln -n 6 –N*, and *bwa samse –n 10000000*. The mapping results were then filtered using *samtools*^6^(v1.9) *view -F 20*. Using the mapping results the clustered UMI pairs were filtered using *gawk* to remove UMI pairs with a coverage < 3x. Potential chimeras were removed by filtering clustered UMI pairs containing sub UMI that was observed in another UMI pair with a higher abundance. The output from these steps was a list of trusted UMI pairs that could be used as references for binning reads.

##### Binning reads according to UMIs

The first 90 bp of each terminal end of the trimmed and filtered reads were extracted with *gawk* and saved into individual files. The UMI pair reference sequences were split into their corresponding sub UMIs and mapped to the read terminals using bwa with the commands: *bwa index, bwa aln -n 3 –N*, and *bwa samse –n 10000000*. The mapping results were then filtered using *samtools view -F 20*. Mapping results from each end of the reads were merged, and a read was binned to a specific UMI pair reference if the following conditions were met: A) Sub UMIs from the same UMI pair were the best hits for both terminal UMIs in the read. B) The mapping difference between the query read and each sub UMI was ≤ 3 bp. C) The mean mapping difference between all of the query reads and the sub UMI was ≤ 3.5 (Nanopore only) or ≤ 3 (PacBio only). D) The ratio between the number of UMI binned reads to the size of the UMI reference cluster was ≤ 10 (Nanopore only). E) The read strand ratio (+/-) was in the interval 10^-0.6^ to 10^0.6^, which is equivalent to the read strand fraction containing the fewest reads comprising at least 25% of the total data amount (Nanopore only). The output from this step was trimmed and filtered reads divided into UMI bins.

##### Generation of UMI consensus sequences

For each individual UMI bin, a consensus sequence was initially generated using *usearch -cluster_fast -id 0.75 -strand both -centroids*, and picking the most abundant centroid. The centroid sequence was used as template for 2 rounds of polishing using all the UMI bin reads with *minimap2*^7^ (v2.17-r954-dirty) *-x map-ont* and *racon*^8^(v1.4.3) *-m 8 -x -6 -g -8 -w 500*.

##### Polishing of UMI consensus sequences (Nanopore data only)

The racon-polished Nanopore consensus sequences were further polished individually by using all of the reads in each UMI bin and two rounds of Medaka (v0.11.2) (https://github.com/nanoporetech/medaka) with the commands *medaka mini_align -m* and *medaka consensus --model r10_min_high_g340 --chunk_len 6000*.

##### Trimming of UMI consensus sequences

The consensus sequences from all UMI bins were then pooled and trimmed and filtered using *cutadapt -m 3000 -M 6000 –g AGRGTTYGATYMTGGCTCAG…GTTTGGCACCTCGATGTCG*. Consensus sequences not containing both primers were discarded.

##### Phasing of consensus sequences and variant calling

Consensus sequences were phased and used to call variants using a custom workflow. The consensus sequences were first filtered to remove any consensus sequences with a read coverage less than the minimum read coverage to obtain >Q40 data quality (25x for ONT UMI, 40x for PB CCS and 3x for PB UMI). The homopolymers were masked in the consensus sequences by converting homopolymers of length ≥3 into length 2 to minimize the effect of homopolymer errors on the phasing accuracy. The masked consensus sequences were dereplicated using *usearch -fastx_uniques -strand both -sizeout -uc* and clustered using two rounds of *usearch -cluster_fast -id 0.995 -strand both -centroids -uc -sort length -sizeout -sizein*, and removing clusters of size < 3. The reads belonging to each cluster were mapped back to the centroid sequence of the cluster using *minimap2 –ax asm5*. Genotype likelihoods were estimated from the mappings with *bcftools* ^9^(v1.9) *mpileup –Ov –d 1000000 –L 1000000 –a “FORMAT/AD,FORMATDP”*, and the results were filtered to show positions of SNPs present in ≥2x coverage using *bcftools view -i ‘AD[0:1-]>2’* for each cluster. The list of SNP positions were used to phase the reads within a cluster, and a variant was called if ≥3 reads supported a combination of SNPs. Consensus reads were then grouped according to called variants, and consensus sequences were re-generated for each variant group. First, the homopolymers were unmasked in the consensus reads and a crude variant-consensus was generated using *usearch -cluster_fast -id 0.99 -strand both -consout –sizeout*. The crude variant-consensus was polished with a workflow using *minimap2 –ax map-ont, bcftools mpileup –Ov –d 1000000 –L 1000000 –a “FORMAT/AD,FORMAT/DP”*, *bcftools norm –Ov, bcftools view -i ‘AD[0:1]/FORMAT/DP>0.5’ –Oz* and *bcftools consensus*.

##### Pipeline parallelization

Many steps in the pipeline have been parallelized using GNU parallel^10^.

#### Data analysis

##### Error profiling

Detection of error was based on a mapping of the sequence data (raw reads, consensus sequences, variant consensus sequences) to curated and non-curated rRNA operon reference sequences from the ZymoBIOMICS Microbial Community DNA Standard (see see *Generation of Reference Sequences for Mock Community*). Mapping was performed with *minimap2 -ax map-ont --cs* and filtered using *samtools view -F 2308*. The references and mappings were imported into the R software environment^11^ (v3.6.0) in RStudio ^12^, where errors in the sequences were profiled using the tidyverse^13^ (v1.2.1) and Biostrings^14^ (v2.52.0) R-packages and custom scripts (see *Resource availability*). In brief, errors and their type (mismatch, deletion, insert) were detected from the SAM --cs tags. The relative positions of the errors was determined with respect to the reference sequence, and this was used to categorize the errors as being within homopolymers regions (hp+) or not (hp-). The error information was combined with metadata from the UMI binning (UMI bin sizes, UMI cluster sizes) and quality analysis (consensus length, UMI bin contamination estimates, ZymoBIOMICS reference-based taxonomy, SILVA taxonomy, chimera detection - see below for details) used to explore and visualize error as a function of such parameters.

##### Taxonomic classification of consensus sequences with mock references

Taxonomic classification of UMI/CCS consensus reads was performed by mapping the reads to curated curated rRNA operon reference sequences from the ZymoBIOMICS Microbial Community DNA Standard with *minimap2 -ax map-ont --cs* and filtered using *samtools view -F 2308*. Read classification was based on best hit.

##### Taxonomic classification of consensus sequences with SILVA database

16S rRNA sequences were extracted from the rRNA operon consensus sequences with *cutadapt--discard-untrimmed -m 1200 -M 2000 -a TGYACWCACCGCCCGTC*. Mapping to the curated ZymoBIOMICS Microbial Community DNA Standard rRNA operon reference sequences and the SILVA 132 SSURef Nr99 database was performed with *minimap2 -ax map-ont --cs* and filtered using *samtools view -F 2308*. Read classification was based on best hit and error rate was calculated as above (see *Error profiling*). The SILVA taxonomy and error rate was used to classify consensus sequences as chimeras or contamination.

##### Estimating UMI bin contamination

Taxonomic classification of raw reads in each UMI bin was performed by mapping the reads to curated rRNA operon reference sequences from the ZymoBIOMICS Microbial Community DNA Standard with *minimap2 -ax map-ont --cs* and filtered using *samtools view -F 2308*. Read classification was based on best hit. Contamination was estimated by calculating the fraction of reads not assigned to the most abundant taxonomy in each UMI bin.

##### Chimera detection

Chimeras in the rRNA operon consensus sequences were detected by *usearch -uchime2_ref-strand plus -mode sensitive,* using our curated rRNA operon reference sequences from the ZymoBIOMICS Microbial Community DNA Standard. Flagged chimeras were manually verified by investigating their error profiles in R (see *Error profiling,* **Supplementary Figure 2**).

##### Examination of relative abundance inconsistencies

We observed a difference between the relative abundance estimated with our UMI consensus data and the theoretical abundance for the rRNA operons of the mock community as reported by the manufacturer. A potential cause of this discrepancy could be due to problems with the DNA composition of the mock community. This was investigated by comparing rRNA operon relative abundance with theoretical relative abundance estimated from metagenomic Nanopore data from ZymoBIOMICS Microbial Community DNA Standard (see *Oxford Nanopore sequencing of mock metagenomic DNA*). Metagenome read lengths were calculated and taxonomy of metagenome reads were classified as above (see *Taxonomic classification of consensus sequences with mock references*) (**Figure S12**). Reference genome size and number of rRNA operons was obtained from the ZymoBIOMICS Microbial Community DNA Standard product manual (_d6305_d6306_zymobiomics_microbial_community_dna_standard.pdf, Ver. 1.1.5). The metagenome data along with the consensus rRNA operon data was imported into the R software environment, and analysed using the tidyverse and Biostrings R-packages along with custom scripts (see *Resource availability*). In short the relative abundance of the consensus rRNA operon data was calculated: Loading…. To calculate the theoretical relative abundance of the rRNA operons using the metagenome data, the metagenome data was first filtered to remove reads < 5000 bp. Read length reflects the DNA template length present in a DNA sample, and < 5000 bp templates are unlikely to contain a complete rRNA operon that can be amplified by PCR, and should therefore be disregarded in an analysis of rRNA operon relative abundance. First the theoretical number of rRNA operons was estimated per reference in the metagenome: Loading…. Then the relative abundance was calculated as above.

##### Analysis of genomic relative abundance and coverage skew due to growth

A bias in relative abundance could occur due to the mock species being in different growth phases at the time of sampling. To investigate the potential contribution of growth to coverage bias, we used metagenomic Nanopore data from ZymoBIOMICS Microbial Community Standards generated internally (see *Oxford Nanopore sequencing of mock metagenomic DNA*) and externally (see *Generation of rRNA operon reference sequences for mock microbial community*). Nanopore data was mapped to each species reference genome using *minimap2 -ax map-ont* and calculated genome read coverage per position by using *samtools depth -aa*. rRNA operon genome coordinates were predicted by barrnap (v.0.9) (available from: https://github.com/tseemann/barrnap) and species genomes were obtained by de novo assembly (see *Generation of Reference Sequences for Mock Community*). The data was imported into R, and used to create read coverage plots (**Supplementary Figure 11**).

##### Investigation of PCR primer match

A bias in relative abundance can be introduced in the first PCR where the rRNA operon is targeted with region-specific primers. If there are mismatches between primers and template, we would expect a lower annealing/amplification efficiency. Primer/template mismatches were estimated using *ipcress* from the package exonerate (v.2.2) (**Supplementary Table 11**).

### E. coli whole genome sequencing with UMIs

#### Sources of DNA

##### Culturing

Escherichia coli str. K-12 substr. MG1655 (DSM 18039) was procured from DSMZ in 2010 and stored at -80 ^₀^C until use. A culture was grown overnight in 2 × 100 mL LB-media (10 g/L NaCl, 10g/L Tryptone, 5 g/L yeast extract) at 37 ^₀^C. Cells were harvested by centrifugation at 7800 RPM for 10 minutes and washed with 1X PBS buffer and finally resuspended in 1 × PBS.

##### DNA extraction

Genomic DNA was extracted with DNeasy PowerSoil (Qiagen, Netherlands) using standard protocol. The DNA concentration was measured on a Qubit 3.0 fluorometer with the Qubit dsDNA HS assay kit (Thermo Fisher Scientific) and the DNA quality was measured by gel electrophoresis on an Agilent 2200 Tapestation using Genomic Screentapes (Agilent Technologies).

#### DNA Sequence Library Preparation

##### Online protocol

An interactive step-by-step protocol is available at protocols.io: (https://www.protocols.io/private/D92C9DC132B111EA92DD0242AC110005).

##### DNA fragmentation

10 µg genomic DNA was fragmented using g-TUBE (Covaris, USA) centrifuged at 8415xg for 60 seconds. The fragmented DNA was purified using CleanPCR (CleanNA, Netherlands) following the manufacturer’s instructions (CleanPCR, manual revision v1.02) with the exception of an EtOH concentration of 80%, post wash dry time of < 3 minutes and 0.8x bead solution/sample ratio and elution 100 µL. DNA concentration and quality was assessed as described earlier.

##### End-repair and UMI adaptor ligation

An end-repair reaction was prepared containing 7 µL NEBNext End Repair Reaction buffer, 3 µL NEBNext End Prep Enzyme mix (New England Biolabs, USA), 2.5 µg fragmented DNA and adjusted to 50 µL with nuclease free water (Qiagen, Netherlands). The reaction was mixed by pipetting 10 times and incubated for 5 minutes at 20 °C and 5 minutes at 65 °C. The DNA was purified using CleanPCR (CleanNA, Netherlands) and its concentration and quality was assessed as described above. UMI adapters were prepared by mixing 25 µL lu_adp_l_v4 (100 µM), 25 µL lu_adp_s_v4 (100 µM), 25 µL 10x NEB T4 DNA ligase buffer (New England Biolabs, USA) and 175 µL nuclease free water (Qiagen, Netherlands) followed by incubation for 5 minutes at 94 °C and 15 minutes at room temperature. The UMI adapters ligation reaction contained 50 µL Blunt/TA ligation Master mix (New England Biolabs, USA), 20 µL adaptor mix prepared above and 1 µg end-repaired DNA in 80 µL nuclease free water. The reaction was mixed by pipetting 10 times and incubated for 10 minutes at 25 °C. The DNA was purified using CleanPCR (CleanNA, Netherlands) and the DNA concentration and quality was assessed as described above.

##### Amplification of UMI-tagged amplicons

4 ng of the adapter-ligated DNA was used as template for an initial PCR amplification of 8 cycles under the same conditions as in ‘*Amplification of UMI-tagged amplicon*s’. The DNA was purified using CleanPCR (CleanNA, Netherlands) and the DNA concentration and quality was assessed as described above. The PCR amplicon was diluted to 2,000 molecules/µL with nuclease free water and used as template for PCR as described in the ‘*Amplification of UMI-tagged amplicons*’ section.

#### Oxford Nanopore sequencing of *E. coli* whole genome UMI amplicons

Sequencing was performed as described in *’rRNA operon UMI sequencing of mock microbial community’* section with the following exceptions. Basecalling was performed with Guppy v3.3.0 (Oxford Nanopore, England) in GPU mode with following modifications to the standard settings --config dna_r10_450bps_hac.cfg model.

#### Data generation workflow

Data generation was performed as described in the *’rRNA operon UMI sequencing of mock microbial community’* section with the following exceptions. Min/max read length cutoffs 2000bp/15000bp. Adaptor sequences used for locating sub UMIs with cutadapt were:

*-g CAAGCAGAAGACGGCATACGAGAT…ACGTGTGCTCTTCCGATCT*

*-G AGATCGGAAGAGCACACGT…ATCTCGTATGCCGTCTTCTGCTTG*.

Only a single round of medaka (v0.8.1) polishing was performed with the commands *medaka mini_align -m* and *medaka consensus --model r10_min_high --chunk_len 6000.* No variants were called.

#### Data analysis

Data analysis was performed as described in *’rRNA operon UMI sequencing of mock microbial community’* with the exception that genomic sequences *Escherichia coli* str_K12_MG1655 genome (NCBI: U00096.3) were used as references when profiling errors (**Supplementary Figure 3**).

### Application of PacBio UMI sequencing of rRNA operons of American Gut

#### Project samples

##### PacBio UMI data generation and processing

PacBio library preparation, sequencing and data generation was performed as described in *’rRNA operon UMI sequencing of mock microbial community’* with the PacBio settings and the following exceptions. 1-2 ng of sample DNA was used as input for ‘*Tagging target gene with UMIs*’. In the third PCR in the library preparation, individual libraries were barcoded by swapping the normal amplification primers for tailed barcode primers (see **Supplementary Table 1**). 25 barcoded libraries were pooled and sent for Sequel II sequencing. After data generation UMI consensus sequences, the data was demultiplexed based on barcodes (see *Resource Availability*).

##### Taxonomic consistency between 16S V4 fragments and full length 16S

To test the consistency of the derived data to the existing Earth Microbiome Project^15^ 16S V4 data, we first extracted full length 16S sequences from the operons using RNAmmer^16^. The sequences were then dereplicated and clustered *de novo* at 99% similarity using VSEARCH 2.7.0^17^using the QIIME 2 version 2019.10^18^ q2-vsearch plugin (parameters: --p-perc-identity 0.99). Taxonomy was assigned against Greengenes 13_5^19^ and the “classify-consensus-vsearch” method of q2-feature-classifier (parameters: --p-strand plus --p-query-cov 0.9 --p-perc-identity 0.9). Next, using redbiom^20^ we obtained EMP 16S V4 Deblur sOTU profiles^21^ for the samples corresponding to the same extracted DNA from Qiita^22^. Both tables were then aggregated to genus level relative abundance, and filtered to only the set of genera in common (n=82) between the two tables. The relative abundance of each genus, per sample, from the full length 16S and the 16S V4 data were then plotted (**Supplementary Figure 14**). Plotting was performed in matplotlib^23^, and Pearson and Spearman correlations were computed using SciPy^24^.

##### Taxonomic specificity of operons from real samples

Sequences of individual rRNA genes were identified from the full-length operon sequences using RNAmmer 1.2 under the “Bacteria” mode. The 16S and 23S rRNA sequences were concatenated with a linker of 20 “N” characters in between. Taxonomic assignment was performed by using the BLASTn algorithm as implemented in NCBI BLAST+ 2.7.1 to align query sequences against the extended “Web of Life” database^25^, which contains all 86,200 non-redundant bacterial and archaeal genomes from NCBI RefSeq and GenBank as of March 2017. The E-value threshold was set as 1e-5, whereas other thresholds were left as default. For each query sequence, hits with a bit score no more than 10% lower than the top hit were selected, and the lowest common ancestor (LCA) of these hits in the taxonomy tree was assigned to the query sequence. This behavior and threshold are consistent with DIAMOND’s taxonomic assignment functionality^26^. The percentage of query sequences assigned to any taxonomic unit at each of the eight standard taxonomic levels were calculated. The taxonomic assignment ratios at species or strain were compared using Pearson’s Chi-square test, as implemented in the “chi2_contingency” function of SciPy 1.3.1.

### Generation of rRNA operon reference sequences for mock microbial community

We obtained raw fast5 files (ENA accession: ERR2887847) from a previously-reported^27^ sequencing effort of the ZymoBIOMICS Microbial Community Standard (D6300, batch ZRC190633) using Oxford Nanopore sequencing. The raw fast5 data was basecalled using the GPU-basecaller guppy v. 2.2.3 with “flipflop” mode. The basecalled reads were mapped to existing reference sequences (updated September 29, 2017; https://s3.amazonaws.com/zymo-files/BioPool/ZymoBIOMICS.STD.refseq.v2.zip) using minimap2 (v.2.12) with default settings. The mapped reads were assembled separately for each reference using minimap2 to create overlaps and miniasm (v.0.3) to perform the assembly with default settings. Raw reads were then mapped to the assembled genomes using minimap2 with default settings and racon (v.1.3.1) was used to construct corrected consensus sequences using default settings. The corrected sequences were subsequently polished with medaka (v.0.6.0) with the “r941_flip_model” model. rRNA operons were extracted from the draft reference genome assemblies using *in silico* PCR with our forward and reverse primers using *ipcress*, and were verified with genome coordinates for rRNA operons predicted by barrnap (**Supplementary Table 11**).

To remove any residual errors from the rRNA operon reference sequences after assembly and polishing, high-quality short reads generated from Illumina sequencing were downloaded from NCBI for each bacterial strain in the mock community (ENA accessions: ERR2935851, ERR2935850, ERR2935852, ERR2935857, ERR2935854, ERR2935853, ERR2935848, ERR2935849) and used for final polishing. The Illumina reads were randomly subsampled to an expected average coverage of 100 for each bacterial strain using the *sample* command in seqtk (v.1.0) (available from: https://github.com/lh3/seqtk). The subsampled Illumina reads were mapped to the draft rRNA operon sequences using minimap2 with the settings: *-ax sr*. The BAM files were sorted and indexed by samtools. We performed variant calling using bcftools (v1.9) with the commands *mpileup* and *call* using the settings: *ploidy =1*. Variant calls were filtered using bcftools *filter* with the settings: *quality > 200*. Variant calls were manually inspected and corrected, if needed, by visualizing mapping profiles in CLC Workbench. Polished consensus sequences were generated with bcftools *consensus* to generate high-quality references (zymo-ref-uniq_2019-10-28.fa) for use in benchmarking error rates in this study (see *Resource availability*). Intragenomic rRNA operons differed by between 0 to 380 bp for the polished rRNA reference sequences (**Supplementary Table 9**).

### Oxford Nanopore sequencing of mock metagenomic DNA

#### Source of DNA

The same ZymoBIOMICS Microbial Community DNA Standard (D6306, lot no. ZRC190811) as described before was used.

#### Oxford Nanopore metagenome sequencing

1500 ng of the mock DNA was used as template for library preparation using the protocol “Genomic DNA by Ligation (SQK-LSK109)” (Oxford Nanopore, England). A MinION R10 flowcell (FLO-MIN110) was used for sequencing on a MinION and MinION software v19.10.1 (Oxford Nanopore, England). Basecalling was performed with Guppy v3.2.2 (Oxford Nanopore, England) in GPU mode with following modifications to the standard settings *--config dna_r10_450bps_hac.cfg*.

### Resources availability

#### Protocols

Interactive protocols are available at protocols.io for the primer based longread UMI approach (https://www.protocols.io/private/F5C5FE21305911EAAC0B0242AC110003) and for the ligation based longread UMI approach https://www.protocols.io/private/D92C9DC132B111EA92DD0242AC110005

#### Code

Source code and analysis scripts are freely available at https://github.com/SorenKarst/longread-UMI-pipeline

#### Data

Raw and assembled sequencing data is available at the European Nucleotide Archive (https://www.ebi.ac.uk/ena) under the project number PRJEB32674 and a complete data overview can be found in **Supplementary Table 12**.

## Supplementary information

**Figure S1:**
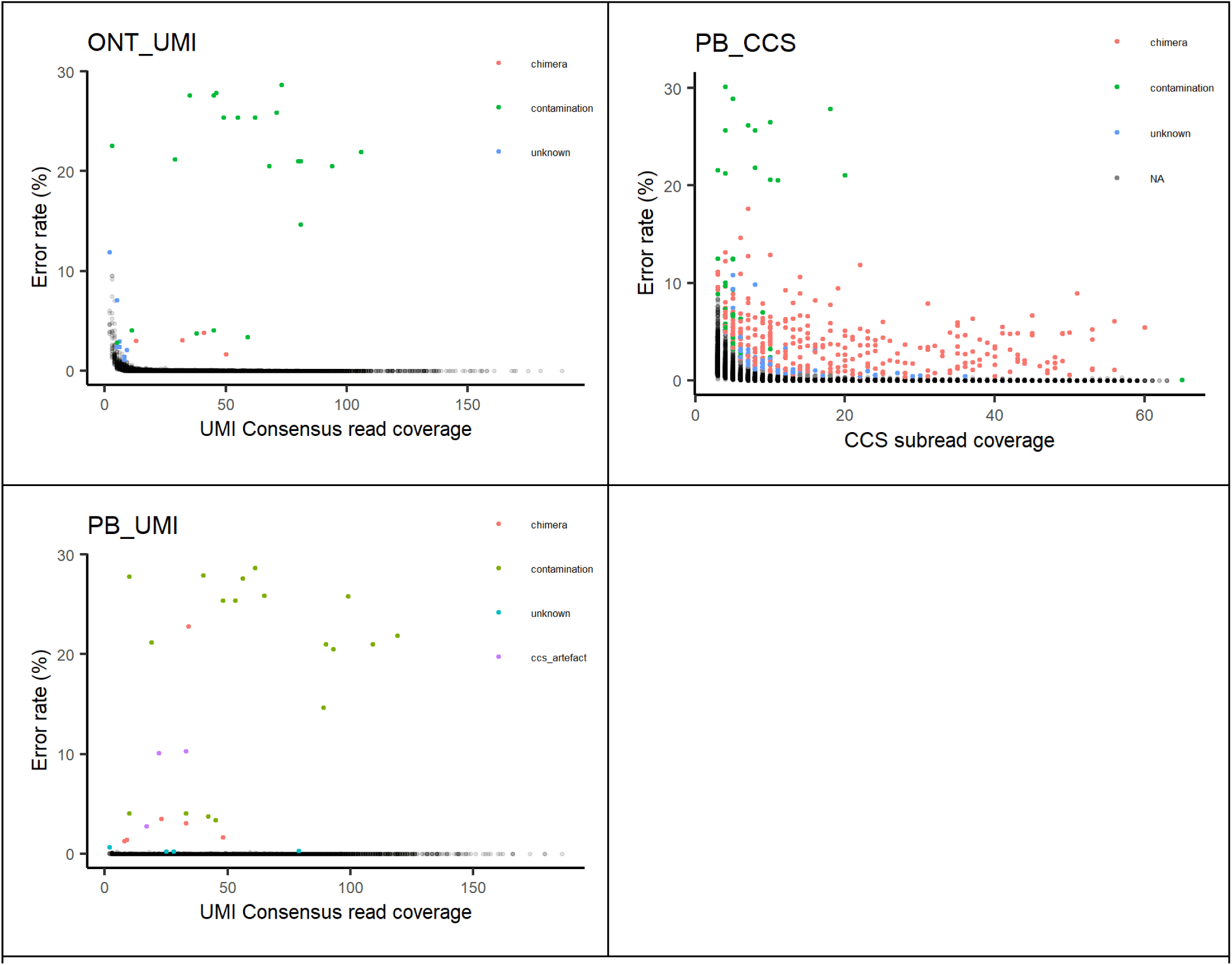
Unfiltered consensus error as a function of read coverage. The plots show consensus error rate as a function of the read coverage before filtering of contamination, chimeras and artefacts. The mean error rate and variance within sliding windows was used to define an error cut-off for that region. Data below the cutoff was flagged as normal (•), and all data above the cut-off was manually inspected and flagged as either chimeric (•), contamination (•), CCS artefact (•) and unknown (•), **see Figure S2**. Contamination originates from PCR reagents and was removed from the data. Chimeras and CCS artefacts were removed from the data and reported in the chimera rate. The CCS artefacts manifested themselves as long stretches of homopolymer inserts, which seem to be present in some of the raw reads and carried over through CCS processing and polishing. Unknown sequences were left in the dataset. The filtered data is presented in Figure 2 in the main article and was used to calculate error statistics. The CCS data shown has been randomly subset to 1/100 (17948 sequences) to make processing and plotting feasible.

**Figure S2:**
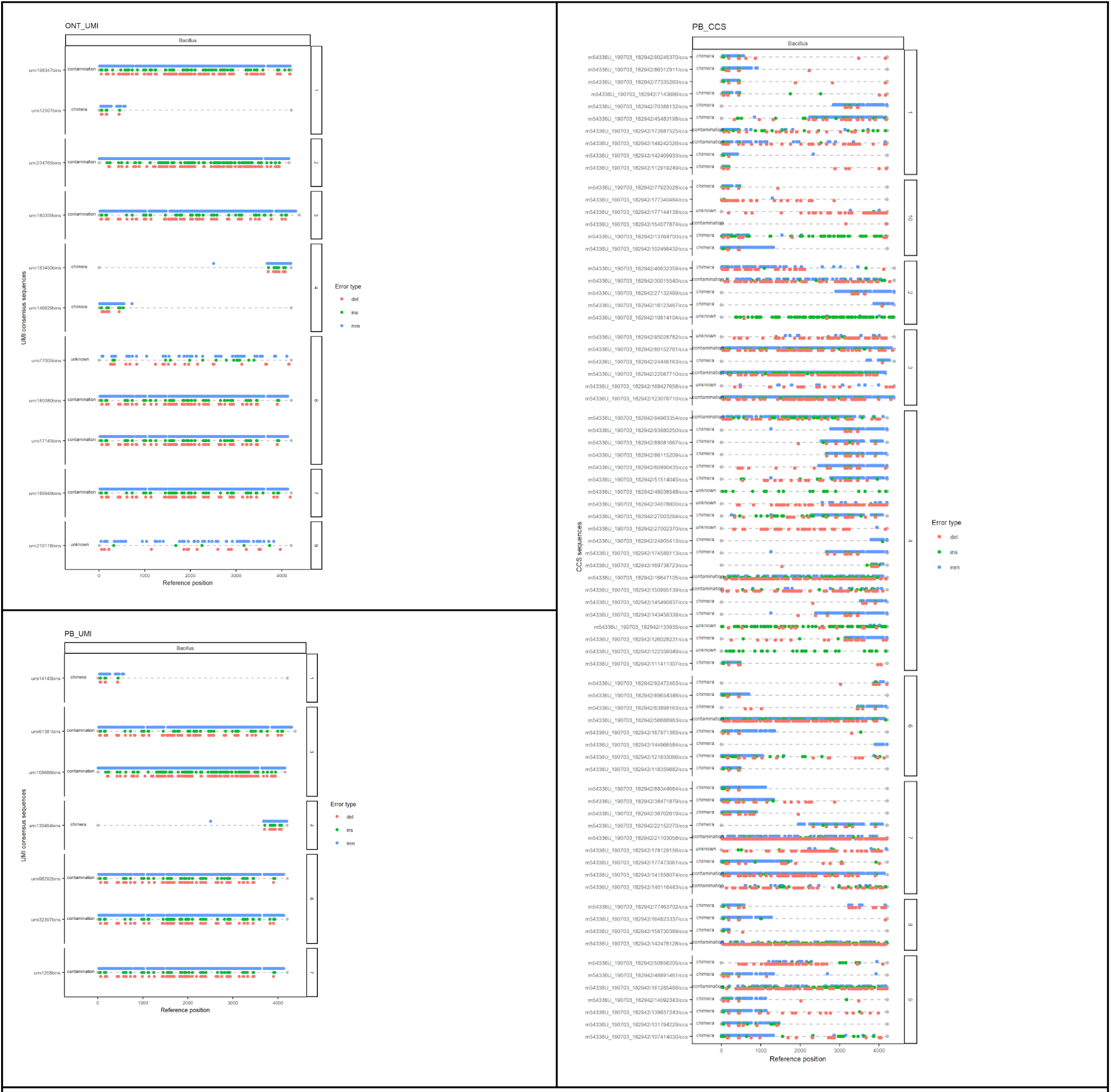
Example of manual inspection of flagged consensus sequences from *Bacillus subtilis*. Outlier consensus sequences are shown for the *Bacillus subtilis* reference. The data is divided by intragenomic operons, and dots signify errors annotated as mismatch, deletion or insert by color. Manual annotations can be seen to the left of the sequences. Chimeras, contamination and artefacts could not be reliably detected by software alone. Therefore, the outliers were flagged depending on error rate and with uchime2_ref chimera detection, and were manually inspected and curated: sequences with errors concentrated in one part of the sequence were flagged as chimeras, sequences with many errors and with a better hit in the SILVA database compared to the ZymoBIOMICS reference were flagged as contamination and sequences with long homopolymer inserts in the PacBio data were flagged as CCS artefacts.

**Figure S3:**
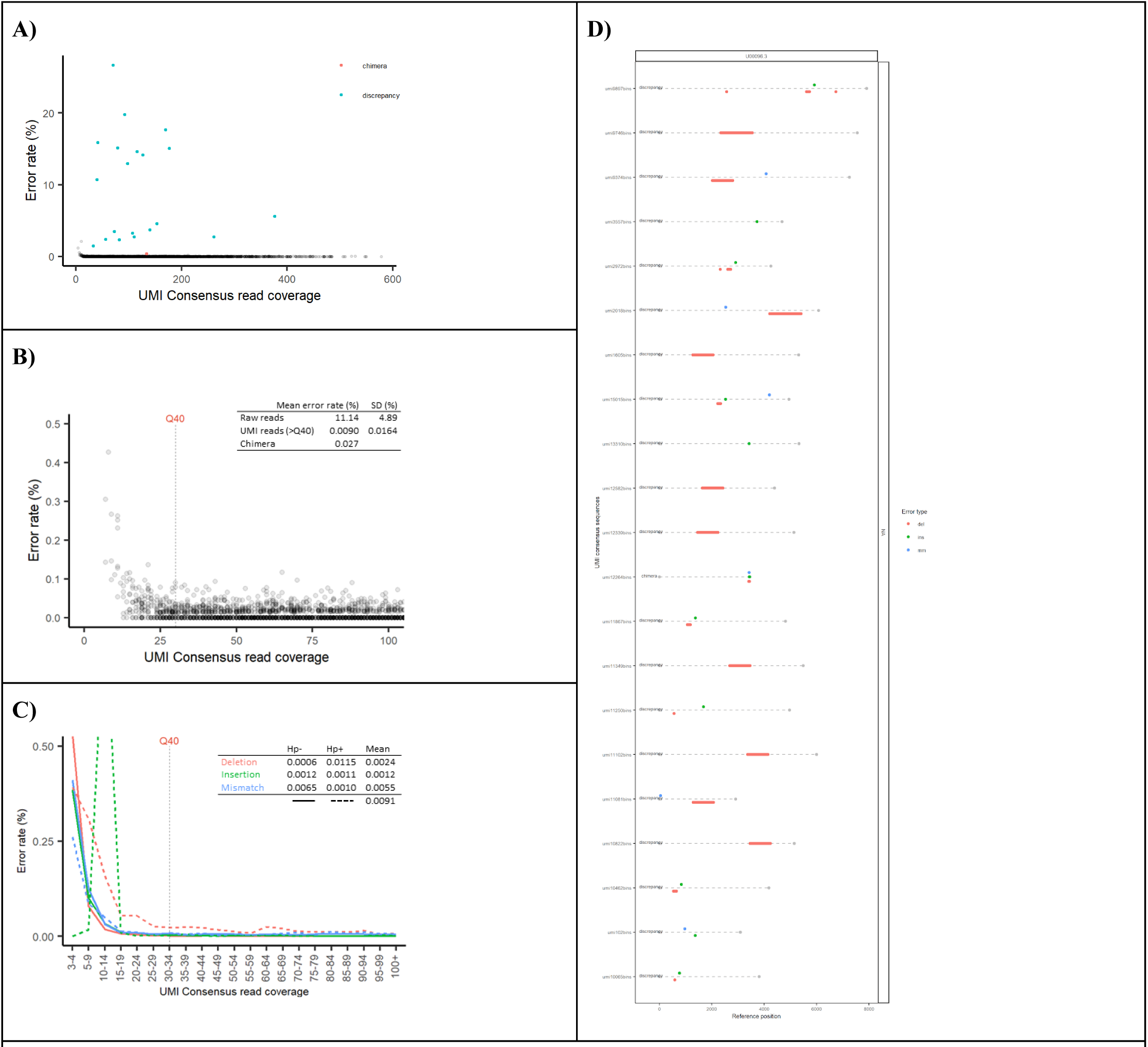
Statistics for ONT UMI consensus sequences from the Escherichia coli genomic shotgun library. We used a shotgun genome library from *E. coli* str. K-12 substr. MG1655 as a proof of concept for using a UMI approach in context of extreme sequence heterogeneity. UMI adaptors were ligated to sheared *E. coli* genomic DNA (mean fragment length ∼8 kbp) and otherwise processed similarly to the amplicon data, generating 3,658 UMI sequences with a read coverage of ≥ 30x with a mean length of 4,476 bp (min = 2000, max = 10578) and a mean error rate of 0.009% and 0.024% chimera rate. A) Error rate of unfiltered consensus sequences versus read coverage. B) Error rate of filtered consensus sequences versus read coverage. C) Error rate divided by type as a function of read coverage and table of error type statistics for data >Q40. D) Error positions and types of flagged outliers.

**Figure S4:**
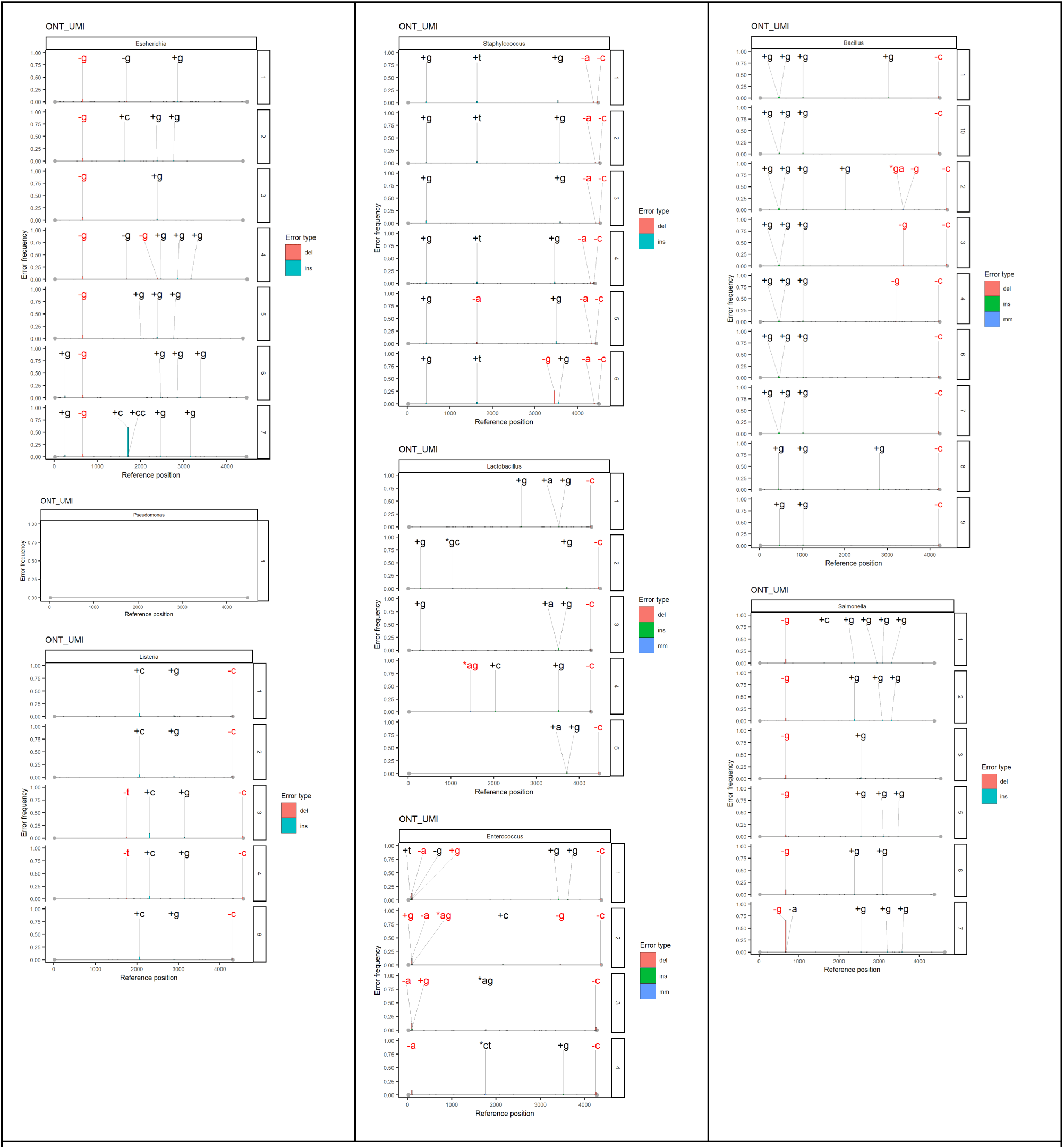
Frequency and reference position of errors in filtered ONT UMI data. Frequency of specific errors are plotted as a function of operon position in bp. The data is divided according to bacterial reference and intragenomic operon number. The error frequency is normalized as fractions of sequences containing the error in that position. Errors with >= 0.01 frequency have been colored by error type and annotated with base-change. +[actg] means insertion -[actg] means deletion and *[actg][actg] means mismatch. Annotated errors in black are in non-homopolymer regions and errors in red are in homopolymer regions.

**Figure S5:**
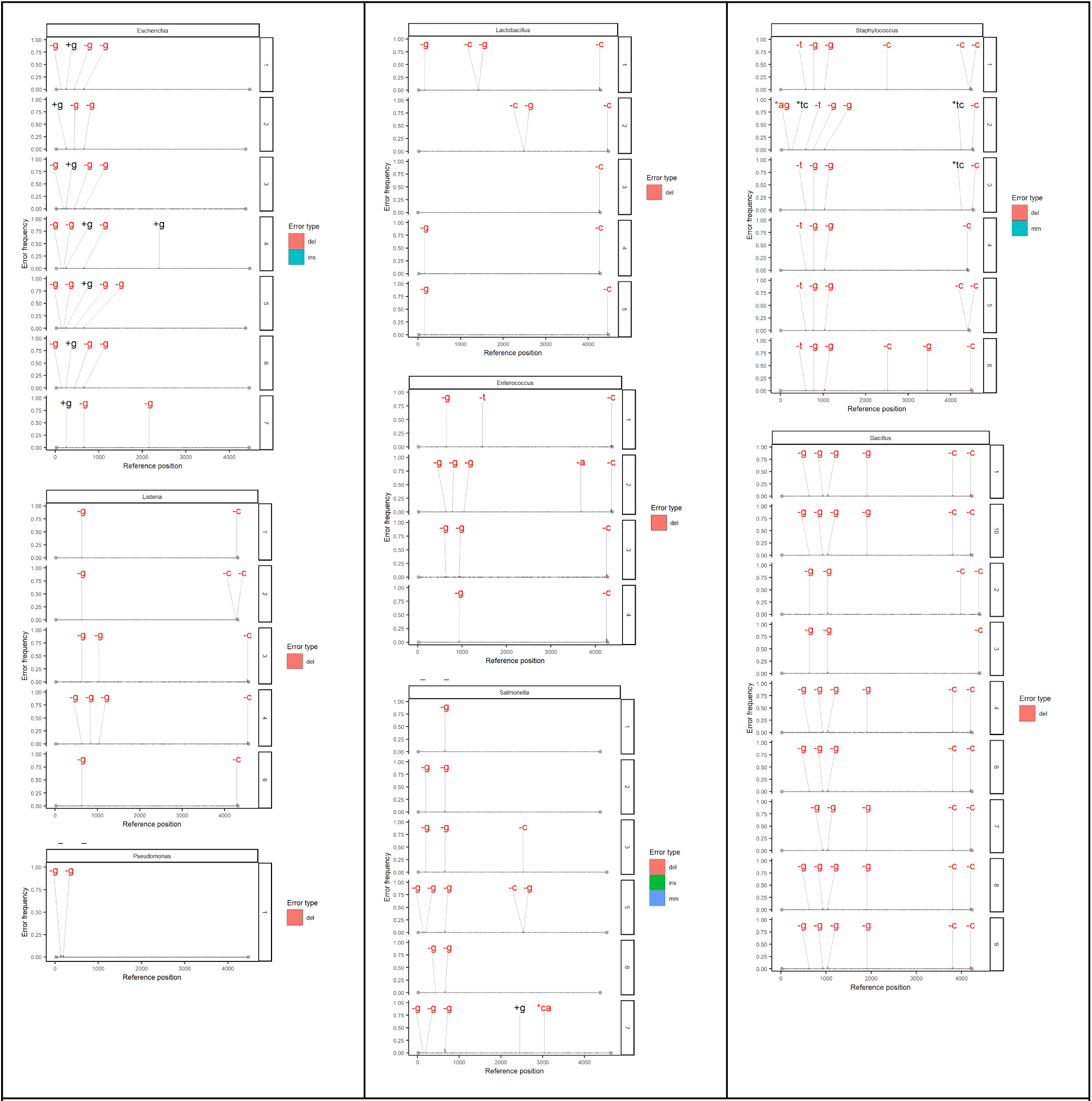
Frequency and reference position of errors in filtered PB CCS data. Frequency of specific errors are plotted as a function of operon position in bp. The data is divided according to bacterial reference and intragenomic operon number. The error frequency is normalized as fractions of sequences containing the error in that position. Errors with >= 0.01 frequency have been colored by error type and annotated with base-change. +[actg] means insertion -[actg] means deletion and *[actg][actg] means mismatch. Annotated errors in black are in non-homopolymer regions and errors in red are in homopolymer regions.

**Figure S6:**
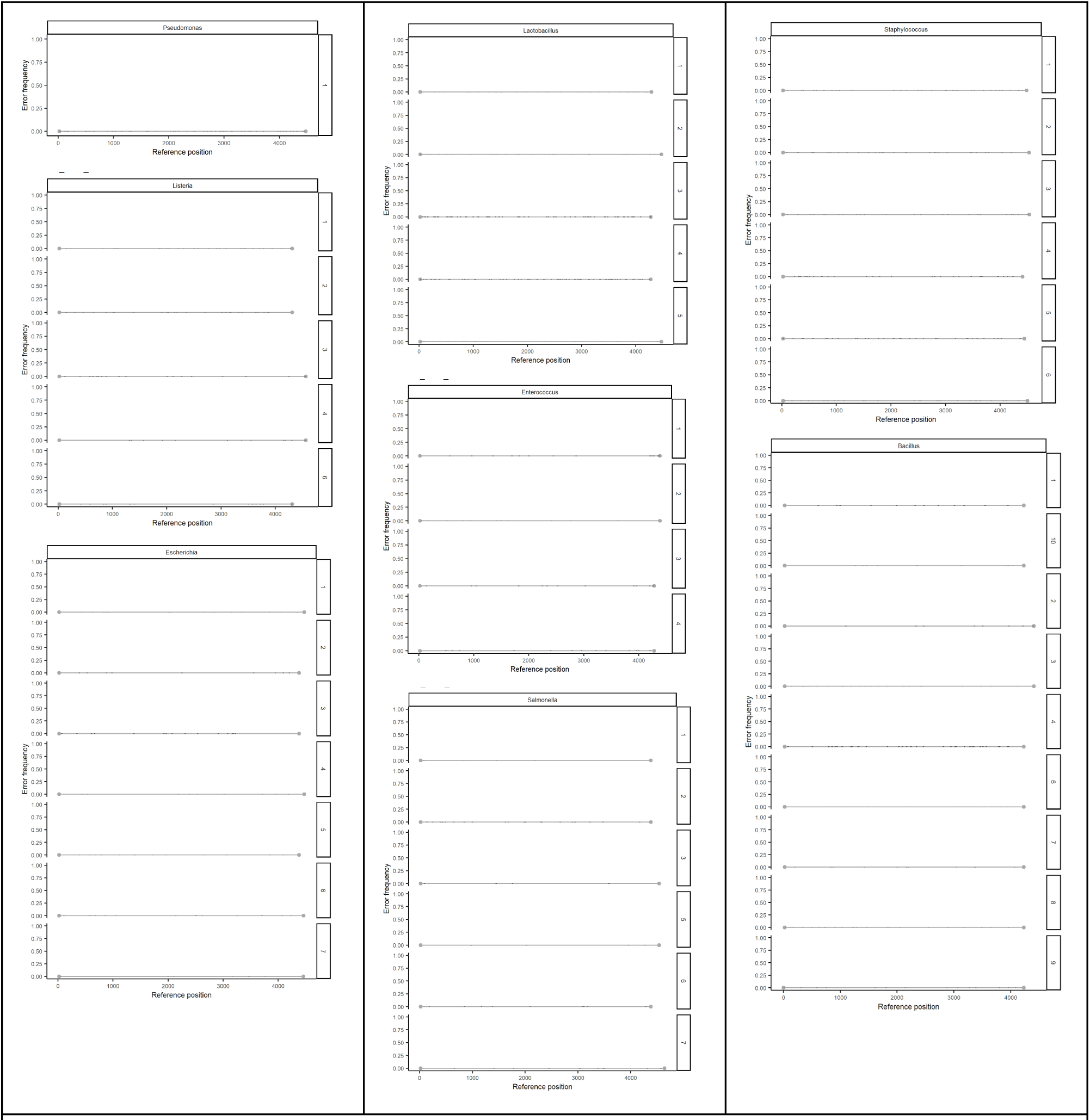
Frequency and reference position of errors in filtered PB UMI data. Frequency of specific errors are plotted as a function of operon position in bp. The data is divided according to bacteria reference and intragenomic operon number. The error frequency is normalized as fractions of sequences containing the error in that position. Errors with >= 0.01 frequency would have been annotated with an error type, but as are no systematic errors and thus there are no annotations.

**Figure S7:**
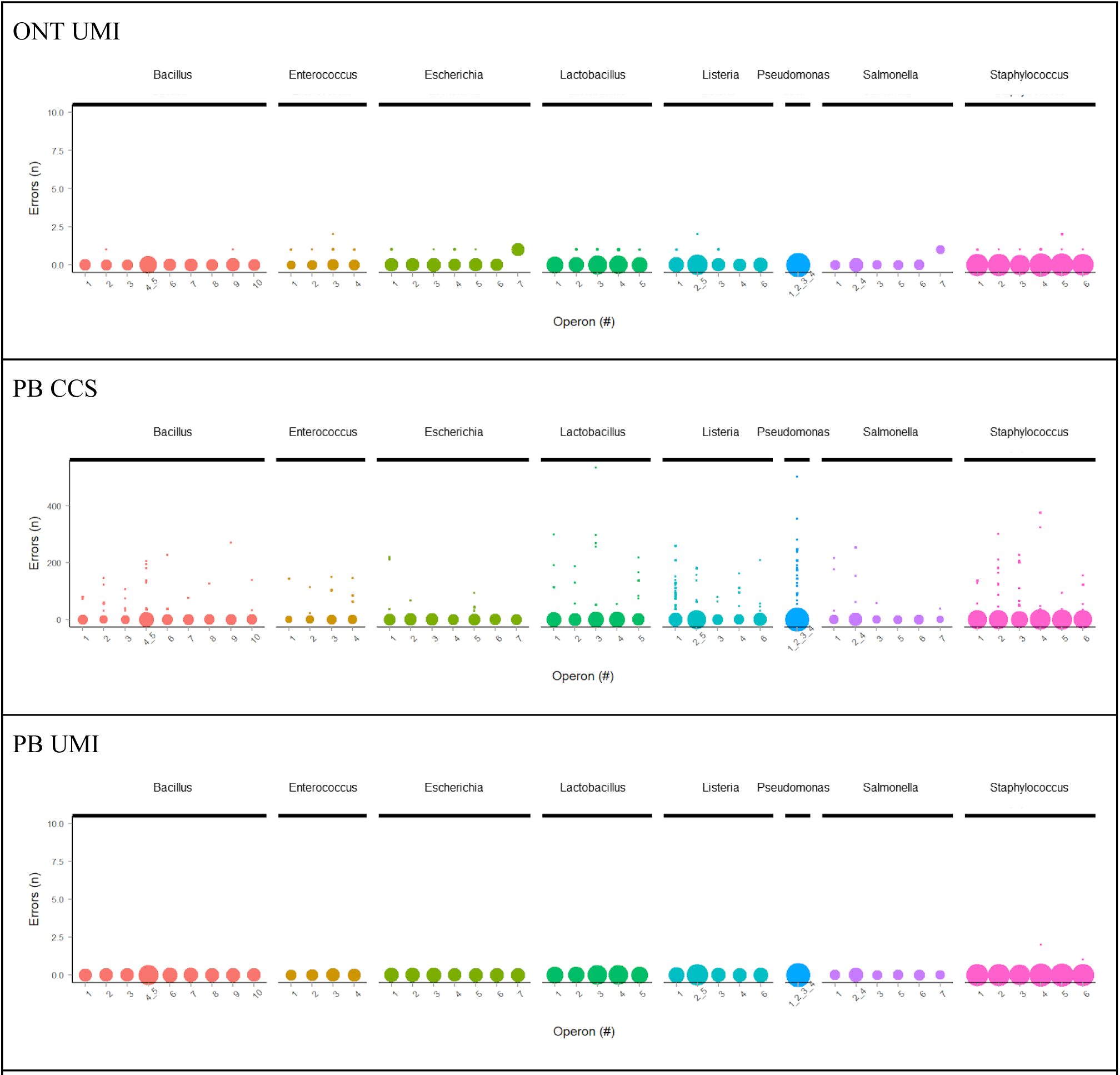
Error count in phased variants. Variant error count as a function of reference intragenomic operon and colored by reference genus. Each point is a variant that is scaled by number of consensus reads used to generate the variant.

**Figure S8:**
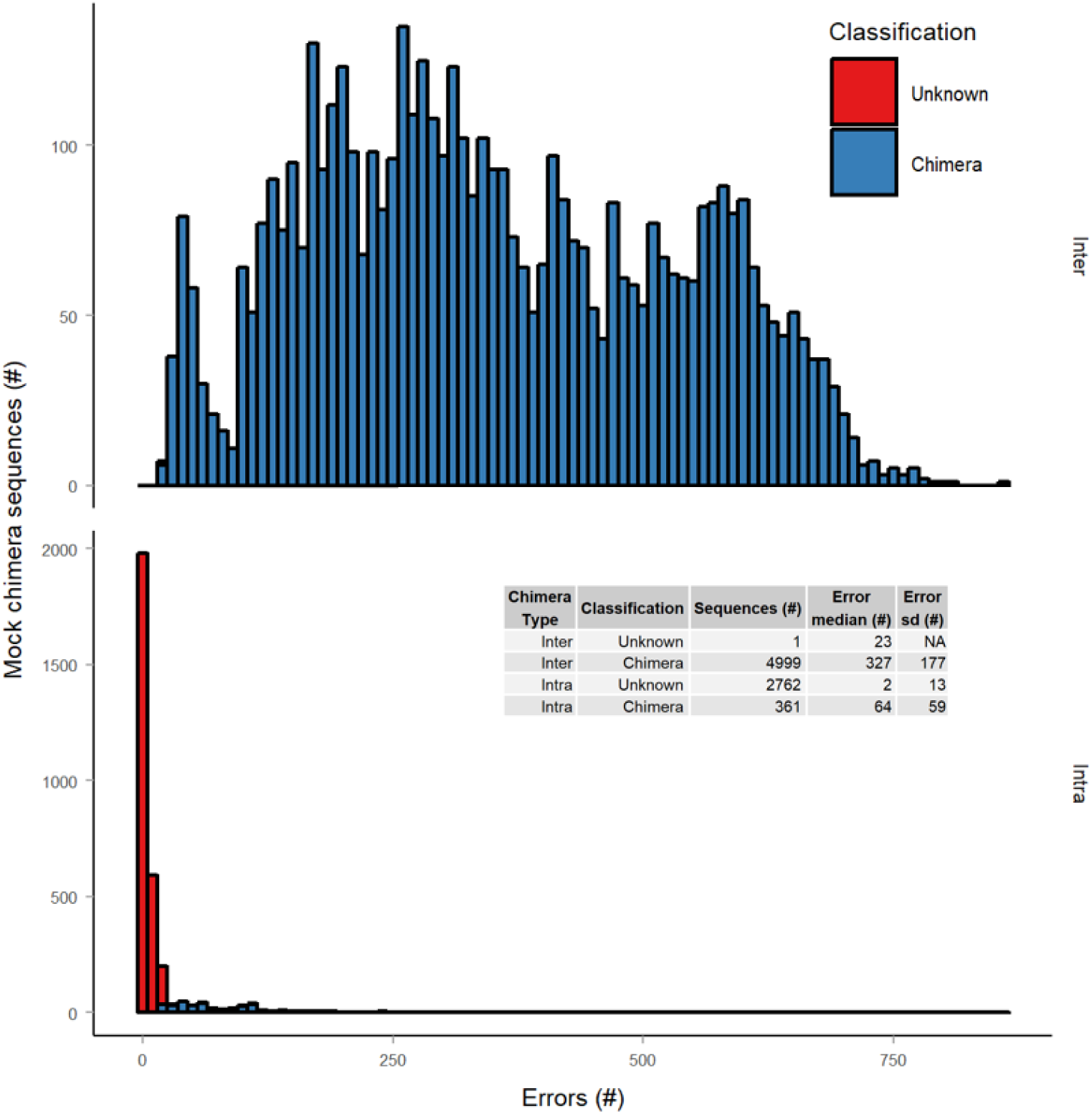
Validation of chimera detection. Chimera detection is notoriously difficult in the presence of sequencing errors and false positives/negatives are unavoidable ^1^. For detecting chimeras in the ZymoBIOMICS Microbial Community DNA Standard rRNA operon data we used uchime2_ref in sensitive mode. To validate that closely related chimeras could be identified with this approach, we generated a chimera dataset from the reference sequences in the mock microbial community, which had between 1 to 842 bp differences to the closest matching references. 99.98% of the inter-species chimeras (n = 5000) were detected along with 11.6% of the intra-species chimeras (n = 3123). The plot shows the test results; the data is divided by inter- and intra-species chimeras, and the x-axis shows the number of differences between the chimera and closest matching reference and the y-axis shows the number of chimeras. It is mainly chimeras with few SNPs that are not classified. The chimera detection method proved to generate false positives when contaminating sequences were present. Hence, detected chimeras were validated by manual inspection (see Figure S2).

**Figure S9:**
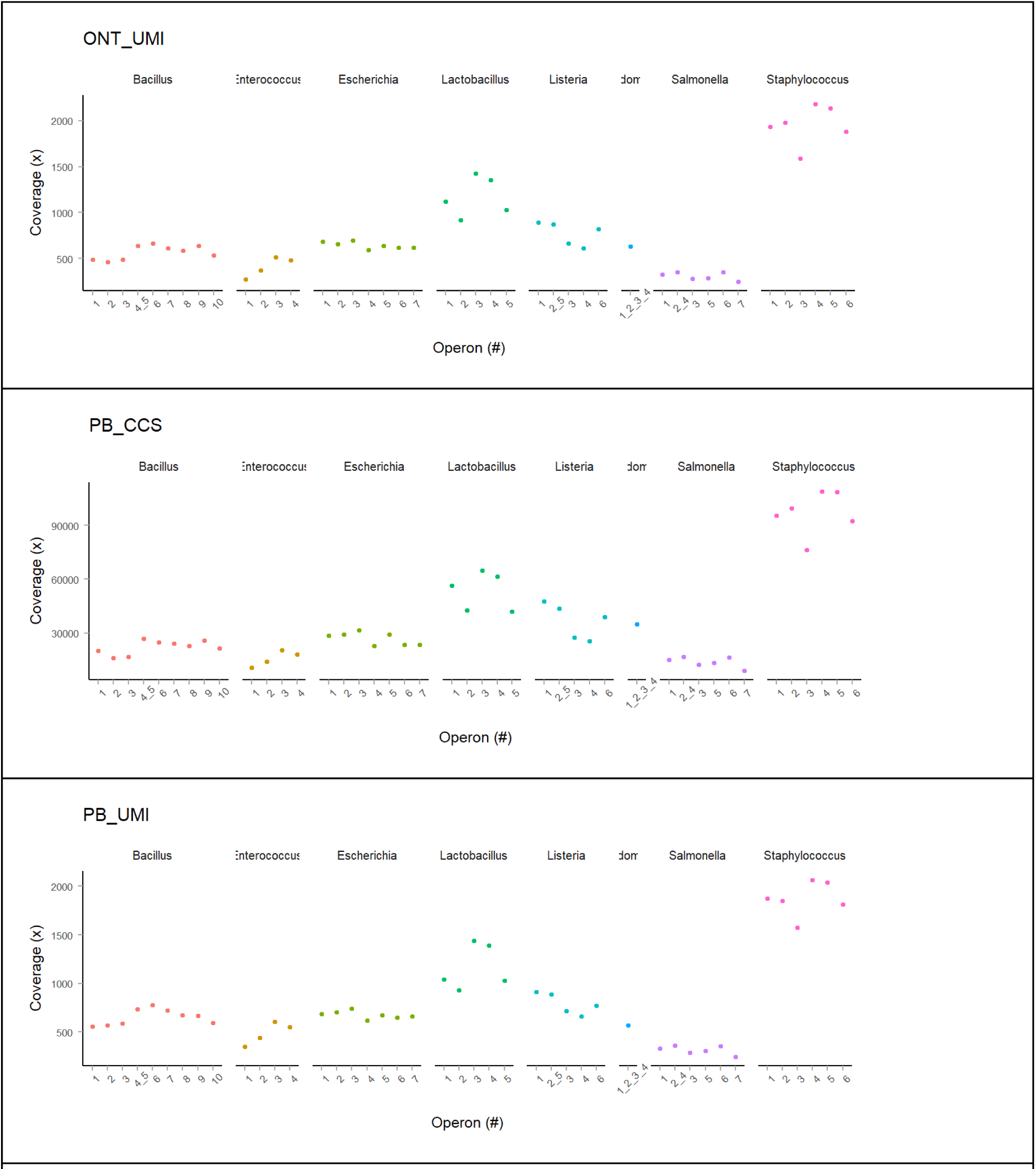
Relative abundance of individual operons. Relative abundance in the form of read coverage as a function of intragenomic operon number for the different reference bacteria. Unfiltered data was used to calculate the relative abundance, as data filtered by read coverage cut-off resulted in adding additional taxa specific biases (see **Figure S10**). The trends were similar in all datasets, irrespective of using UMIs or not.

**Figure S10:**
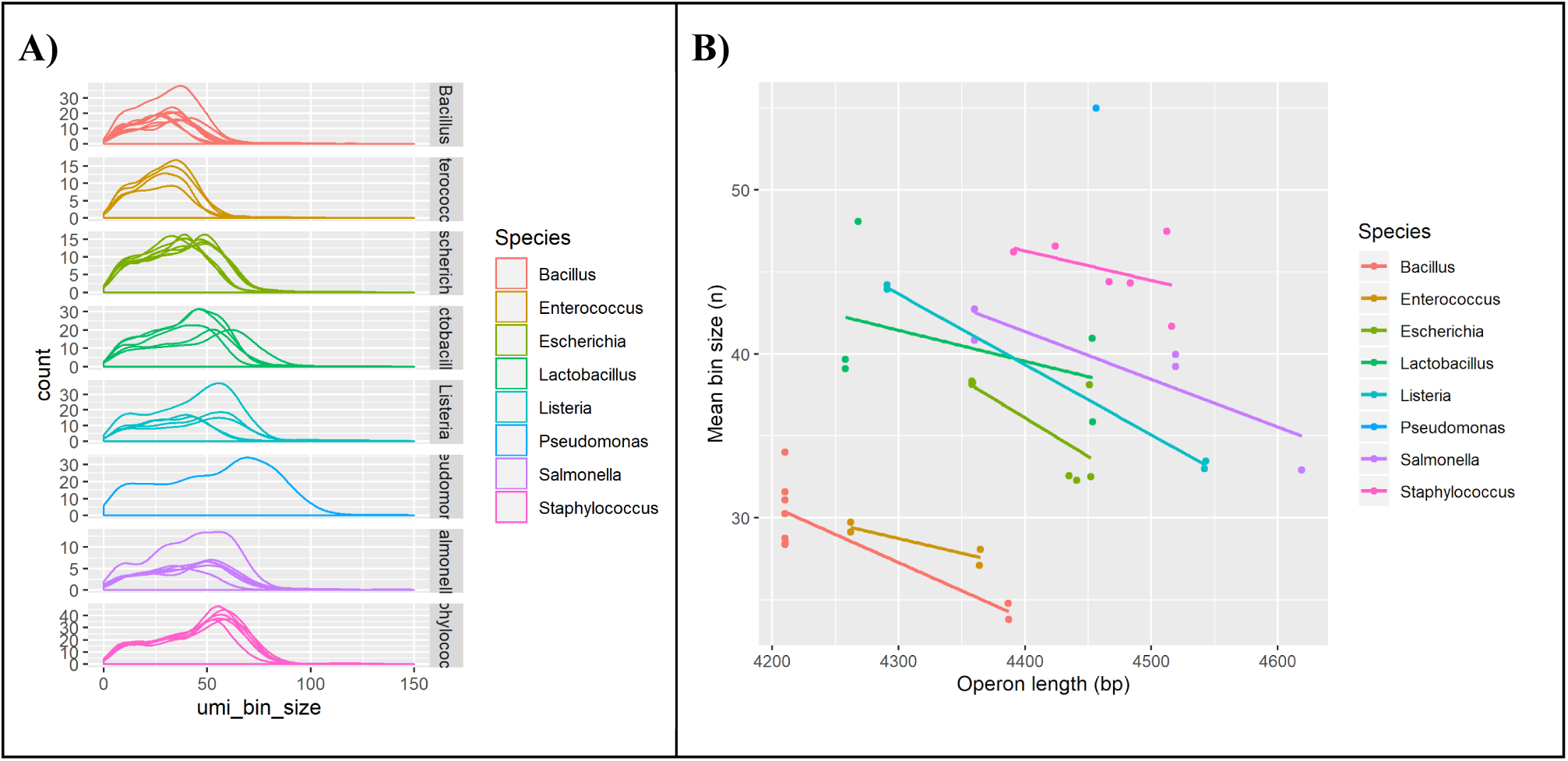
Correlation between operon length and UMI bin size. (A) Density plots showing the number of sequences as function of UMI bin size divided by intragenomic operon number and reference genus. (B) Correlation between mean UMI bin size and rRNA operon length coloured by genus. PB UMI data was used to generate the plots. UMI bin size can be used as a proxy for PCR efficiency. UMI consensus sequences originate from a single molecule, and all tagged molecules in sample are amplified using the same synthetic primers and under the same PCR conditions. Differences in relative read coverage per molecule in the final PCR product should therefore only originate from length and nucleotide composition based PCR efficiency biases. There is a clear trend of efficiency depending on length and taxonomy. However, the UMI approach should mitigate the post UMI tagging PCR biases shown in the above plots, which means the observed overall bias in relative abundance (**Table S5**) must be introduced in the UMI PCR or be present in the template to begin with.

**Figure S11:**
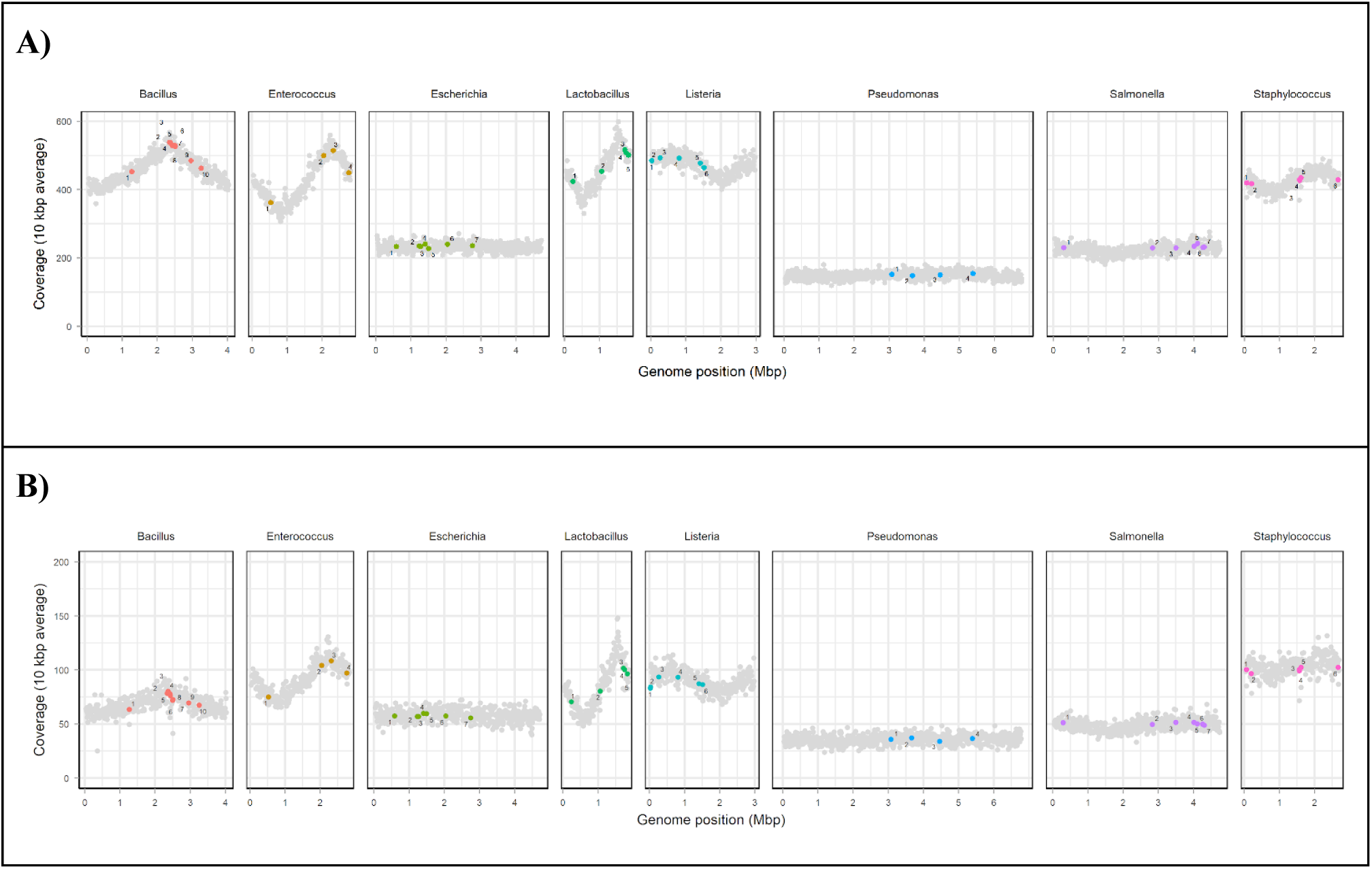
Mock microbial community read coverage across genomes. Read coverage profiles of the ZymoBIOMICS Microbial Community DNA Standard based on shotgun Nanopore sequencing data. Each grey point is the average coverage value of a 10 kbp region within the genome. Colored points represents the position of the individual intragenomic rRNA operons. A) is data generated from a ZymoBIOMICS Microbial Community Standard (even) [product D6300, batch ZRC190633] by the Loman lab ^2^ using the Nanopore GridION and a R9.4.1 flowcell. B) is our data generated from a ZymoBIOMICS Microbial Community DNA Standard (even) [product D6306, batch ZRC190811] using Nanopore MinION and a R10 flowcell.

**Figure S12:**
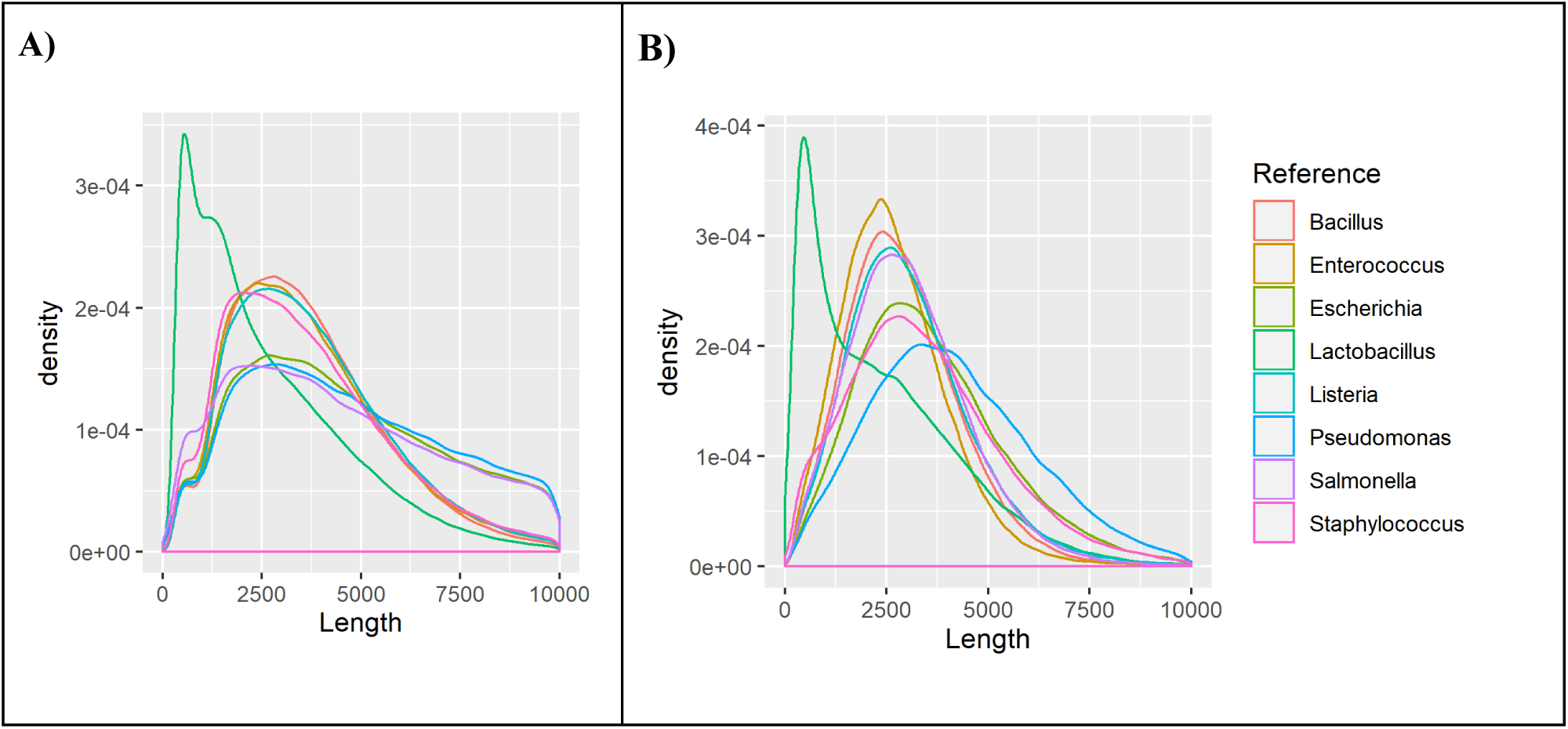
Read size distribution of mock community metagenome data. Each line plot represents the read size distribution from each mock community species estimated from the Nanopore metagenome data. A) is data generated from a ZymoBIOMICS Microbial Community Standard (even) [D6300, batch ZRC190633] by the Loman lab ^2^ using the Nanopore GridION and a R9.4.1 flowcell. B) is in-house data generated from a ZymoBIOMICS Microbial Community DNA Standard (even) [product D6306, batch ZRC190811] using Nanopore MinION and a R10 flowcell. Some species have significantly more high molecular weight DNA over 5000 bp compared to some of the other species, which impacts the effective template availability in PCR for long amplicons. For the Loman lab data, the distinct gram+/- dependent trends fragment length is anticipated from their two step extraction protocol used on the mock community. The in-house generated data is generated from DNA standard prepared by the vendor, and here the taxa trend is not as pronounced, but the effect on the effective relative abundance is still dramatic (**Table S5**). We can only speculate about the cause, but different extraction methods could be the reason. If all samples had been extracted with the same method, the result should be more similar fragment distributions as indicated by the Loman group data. The Nanopore library preparation and sequencing likely had an impact on the observed read fragment distributions, but the Loman group data indicates that extraction method probably is the most important factor.

**Figure S13:**
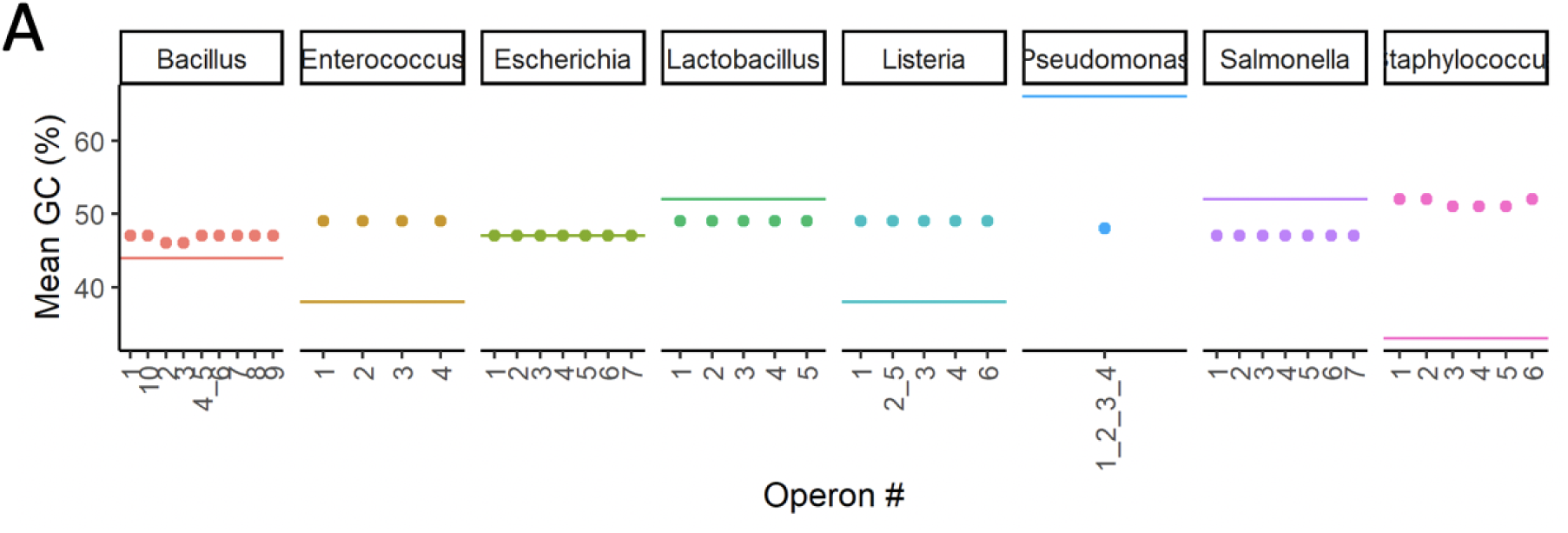
Genome and operon GC content. (A) Mean GC content of the genome (solid lines) along with the GC content of each rRNA operon (points), arranged by species. Despite different average genome content the GC content of the intragenomic operons is very similar across species (46% to 52%), and can be very different from the genome GC content.

**Figure S14.**
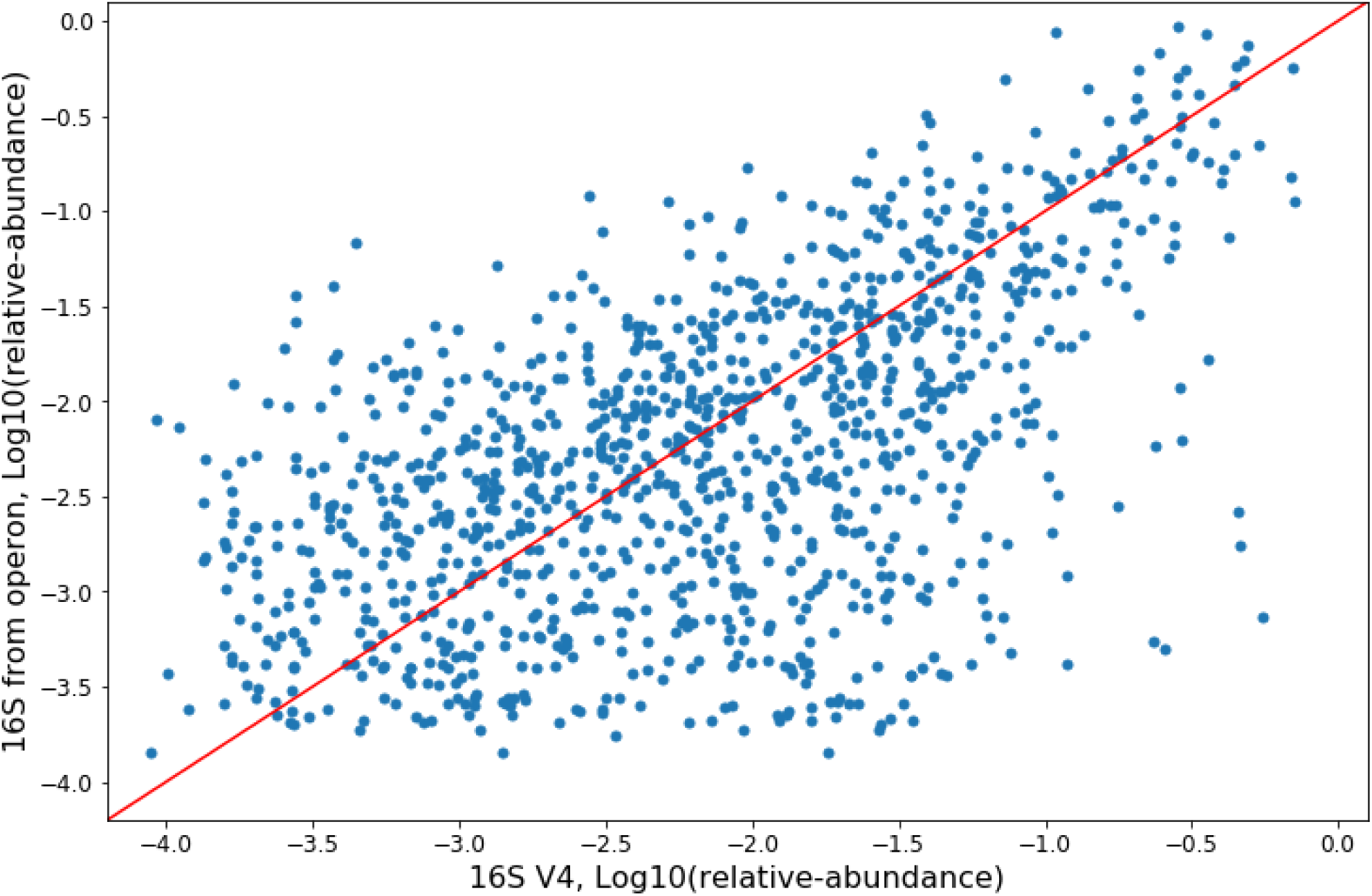
A scatter plot of both molecular preparations showing the relative abundances of bacterial genera within samples (n=70). Each point represents a bacterial genus within a given sample, and shows the observed relative abundance from Earth Microbiome Project 16S V4 derived data, and the observed relative abundance of the full operon data based on taxonomic annotation of the full length 16S. The red line depicts y=x. The relative abundances are significantly correlated with existing V4 data^3^(Spearman: r=0.527, p=1.449e-80; Pearson: r=0.553, p=4.555e-90).

**Figure S15:**
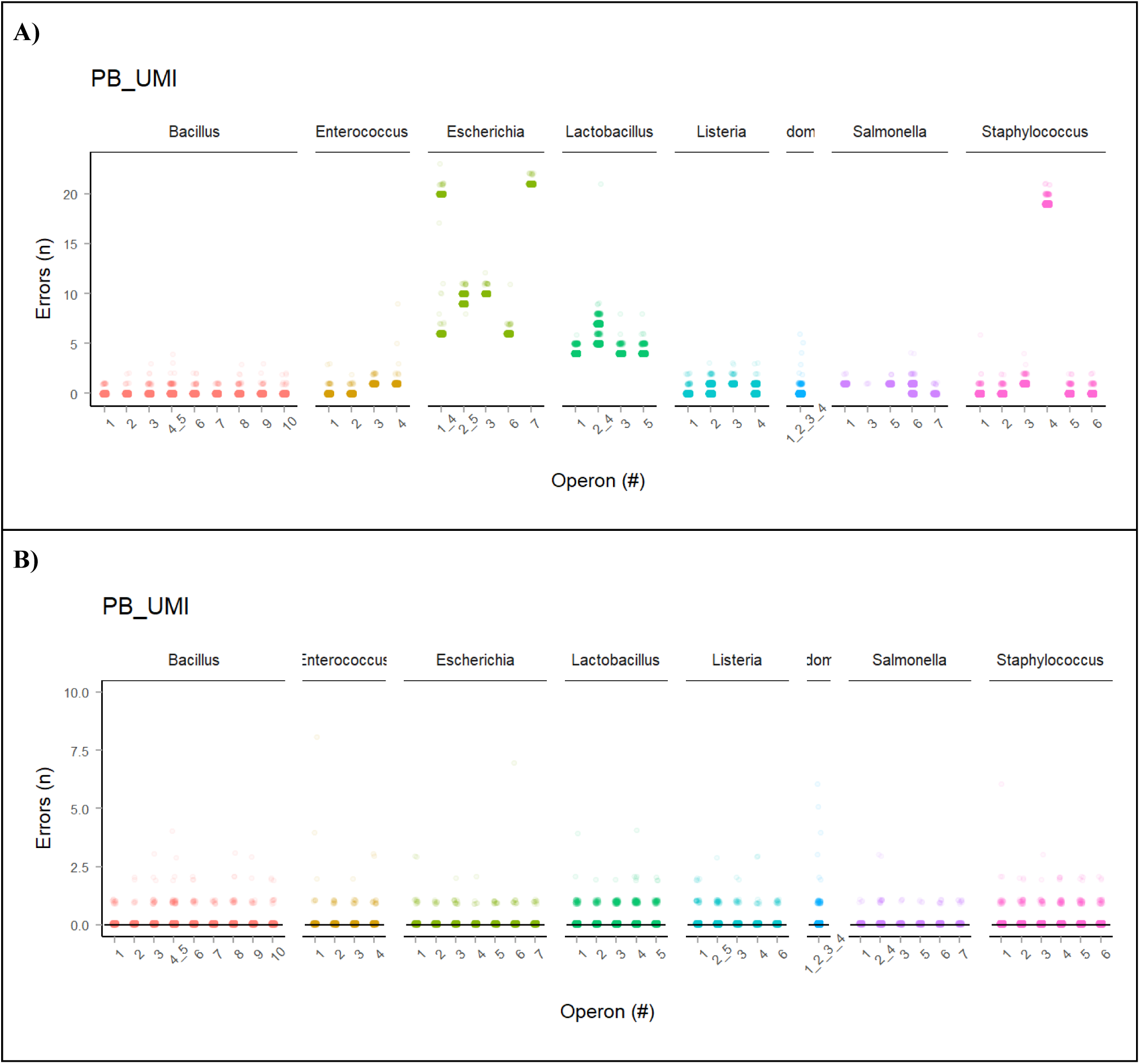
Number of errors in PB UMI consensus sequences using non-curated and curated references. A) The number of errors in the PB UMI consensus sequences estimated based on non-curated rRNA reference sequences (updated September 29, 2017; https://s3.amazonaws.com/zymo-files/BioPool/ZymoBIOMICS.STD.refseq.v2.zip). B) The number of errors in the PB UMI consensus sequences estimated based on curated rRNA reference sequences. Each point represents a UMI consensus sequence that aligns to a specific reference operon. These observations were confirmed with ONT UMI data indicated errors in the available reference genomes, as was also reported by others^4^. To generate improved rRNA operon references, we first used a long-read first assembly approach, in which publicly available ONT sequence data of the Zymo mock community^2^ was assembled into individual reference genomes with Miniasm^5^ followed by Racon and Medaka polishing. rRNA operons were extracted from the high-quality long-read assemblies, and SNPs with no Illumina short-read support were manually curated, which were mainly indel errors in homopolymers. In total, we found 49 bacterial rRNA operons with 4-10 copies/species, where 43 operons were unique and had 1-379 intra-species differences (**Table S9**). The mean difference between the original references and our curated sequences was 0.063% (∼2.8 SNP/operon), with a range of 0 – 0.47% (0 – 21 SNP/operon).

**Table S1:**
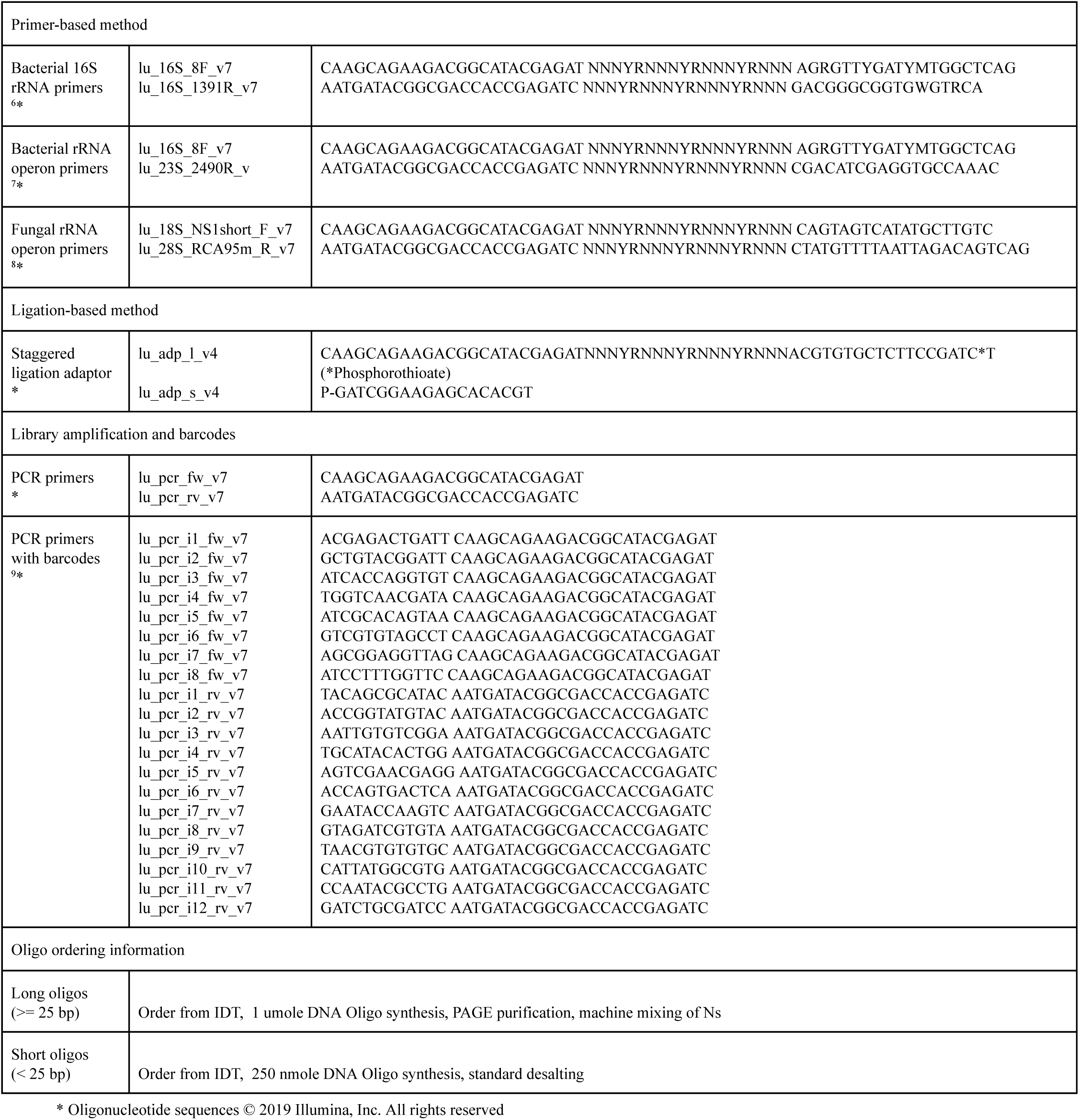
Primers and adaptors used for data generation.

**Table S2:** Consensus error calculated for 5 bp intervals for ZymoBIOMICS Microbial Community DNA Standard rRNA amplicon libraries.

| ONT_UMI |  |  |  |  |  | PB_CCS |  |  |  |  |  |
| --- | --- | --- | --- | --- | --- | --- | --- | --- | --- | --- | --- |
| UMI bin size | Total Error rate (%) | Mismatch Error rate (%) | Insert Error rate (%) | Deletion Error rate (%) | Sequence count (n) | UMI bin size | Total Error rate (%) | Mismatch Error rate (%) | Insert Error rate (%) | Deletion Error rate (%) | Sequence count (n) |
| 2-4 | 2.748 | 1.290 | 0.437 | 1.021 | 87 | 2-4 | 1.591 | 0.290 | 0.690 | 0.611 | 3751 |
| 5-9 | 0.329 | 0.129 | 0.064 | 0.136 | 846 | 5-9 | 0.371 | 0.036 | 0.144 | 0.192 | 4970 |
| 10-14 | 0.071 | 0.018 | 0.022 | 0.031 | 1570 | 10-14 | 0.081 | 0.005 | 0.023 | 0.053 | 2613 |
| 15-19 | 0.029 | 0.006 | 0.011 | 0.012 | 1956 | 15-19 | 0.040 | 0.003 | 0.010 | 0.026 | 1697 |
| 20-24 | 0.015 | 0.002 | 0.006 | 0.006 | 2424 | 20-24 | 0.024 | 0.003 | 0.006 | 0.015 | 1112 |
| 25-29 | 0.010 | 0.002 | 0.005 | 0.004 | 2767 | 25-29 | 0.017 | 0.002 | 0.004 | 0.011 | 844 |
| 30-34 | 0.008 | 0.001 | 0.004 | 0.003 | 2779 | 30-34 | 0.013 | 0.002 | 0.003 | 0.008 | 673 |
| 35-39 | 0.006 | 0.001 | 0.003 | 0.002 | 2769 | 35-39 | 0.012 | 0.002 | 0.004 | 0.006 | 565 |
| 40-44 | 0.005 | 0.001 | 0.002 | 0.002 | 2746 | 40-44 | 0.009 | 0.003 | 0.002 | 0.005 | 480 |
| 45-49 | 0.004 | 0.001 | 0.002 | 0.001 | 2617 | 45-49 | 0.006 | 0.003 | 0.001 | 0.003 | 370 |
| 50-54 | 0.003 | 0.000 | 0.002 | 0.001 | 2254 | 50-54 | 0.009 | 0.004 | 0.001 | 0.004 | 276 |
| 55-59 | 0.003 | 0.001 | 0.002 | 0.001 | 1956 | 55-59 | 0.010 | 0.003 | 0.001 | 0.006 | 175 |
| 60-64 | 0.003 | 0.001 | 0.001 | 0.001 | 1624 | 60-64 | 0.009 | 0.000 | 0.001 | 0.008 | 21 |
| 65-69 | 0.002 | 0.001 | 0.001 | 0.001 | 1318 | 65-69 | NA | NA | NA | NA | 0 |
| 70-74 | 0.002 | 0.000 | 0.001 | 0.001 | 826 | 70-74 | NA | NA | NA | NA | 0 |
| 75-79 | 0.001 | 0.000 | 0.001 | 0.000 | 596 | 75-79 | NA | NA | NA | NA | 0 |
| 80-84 | 0.001 | 0.000 | 0.001 | 0.000 | 395 | 80-84 | NA | NA | NA | NA | 0 |
| 85-89 | 0.001 | 0.000 | 0.000 | 0.000 | 243 | 85-89 | NA | NA | NA | NA | 0 |
| 90-94 | 0.001 | 0.000 | 0.000 | 0.000 | 149 | 90-94 | NA | NA | NA | NA | 0 |
| 95-99 | 0.002 | 0.001 | 0.001 | 0.001 | 103 | 95-99 | NA | NA | NA | NA | 0 |
| 100+ | 0.001 | 0.000 | 0.001 | 0.000 | 215 | 100+ | NA | NA | NA | NA | 0 |

| PB_UMI |  |  |  |  |  |
| --- | --- | --- | --- | --- | --- |
| UMI bin size | Total Error rate (%) | Mismatch Error rate (%) | Insert Error rate (%) | Deletion Error rate (%) | Sequence count (n) |
| 2-4 | 0.007 | 0.000 | 0.003 | 0.003 | 529 |
| 5-9 | 0.001 | 0.001 | 0.000 | 0.000 | 2336 |
| 10-14 | 0.001 | 0.001 | 0.000 | 0.000 | 2312 |
| 15-19 | 0.001 | 0.001 | 0.000 | 0.000 | 2427 |
| 20-24 | 0.001 | 0.001 | 0.000 | 0.000 | 2685 |
| 25-29 | 0.001 | 0.001 | 0.000 | 0.000 | 3032 |
| 30-34 | 0.001 | 0.001 | 0.000 | 0.000 | 3372 |
| 35-39 | 0.000 | 0.000 | 0.000 | 0.000 | 3315 |
| 40-44 | 0.001 | 0.000 | 0.000 | 0.000 | 3353 |
| 45-49 | 0.001 | 0.000 | 0.000 | 0.000 | 3359 |
| 50-54 | 0.001 | 0.000 | 0.000 | 0.000 | 3275 |
| 55-59 | 0.000 | 0.000 | 0.000 | 0.000 | 2922 |
| 60-64 | 0.000 | 0.000 | 0.000 | 0.000 | 2421 |
| 65-69 | 0.000 | 0.000 | 0.000 | 0.000 | 1654 |
| 70-74 | 0.001 | 0.000 | 0.000 | 0.000 | 1032 |
| 75-79 | 0.001 | 0.000 | 0.000 | 0.001 | 600 |
| 80-84 | 0.000 | 0.000 | 0.000 | 0.000 | 352 |
| 85-89 | 0.000 | 0.000 | 0.000 | 0.000 | 205 |
| 90-94 | 0.000 | 0.000 | 0.000 | 0.000 | 137 |
| 95-99 | 0.001 | 0.001 | 0.000 | 0.000 | 99 |
| 100+ | 0.000 | 0.000 | 0.000 | 0.000 | 279 |

**Table S3:** Error rate divided by homopolymer type (nucleotide/length) for ZymoBIOMICS Microbial Community DNA StandardZymo Mock Community rRNA amplicon libraries.

| ONT_UMI |  |  |  |  | PB_CCS |  |  |  |  |
| --- | --- | --- | --- | --- | --- | --- | --- | --- | --- |
| hp_len | A | C | G | T | hp_len | A | C | G | T |
| 3 | 0.008 | 0.003 | 0.003 | 0.001 | 3 | 0.004 | 0.011 | 0.013 | 0.004 |
| 4 | 0.001 | 0.004 | 0.005 | 0 | 4 | 0.01 | 0.066 | 0.045 | 0.016 |
| 5 | 0.001 | 0.342 | 0.005 | 0.02 | 5 | 0.046 | 0.551 | 0.175 | 0.04 |
| 6 | 0.25 | NA | 0.184 | 0.001 | 6 | 0.014 | NA | 0.112 | 0.15 |
| 7 | 0 | NA | 4.401 | NA | 7 | 0.053 | NA | 0.359 | NA |
| 3 | 1492724 | 801057 | 1113259 | 681436 | 3 | 8426922 | 4487576 | 6319282 | 3845627 |
| 4 | 369567 | 204350 | 457313 | 138154 | 4 | 2084166 | 1155005 | 2576527 | 775562 |
| 5 | 70619 | 32762 | 84371 | 19273 | 5 | 407325 | 180944 | 492265 | 101264 |
| 6 | 2934 | 0 | 24239 | 15668 | 6 | 16425 | 0 | 145353 | 86650 |
| 7 | 1155 | 0 | 1305 | 0 | 7 | 6171 | 0 | 6805 | 0 |

| PB_UMI |  |  |  |  |
| --- | --- | --- | --- | --- |
| hp_len | A | C | G | T |
| 3 | 0 | 0.002 | 0.001 | 0 |
| 4 | 0 | 0.002 | 0.002 | 0 |
| 5 | 0 | 0.002 | 0.003 | 0.001 |
| 6 | 0 | NA | 0.003 | 0 |
| 7 | 0 | NA | 0.014 | NA |
| 3 | 2521020 | 1376806 | 1897615 | 1140578 |
| 4 | 618398 | 349152 | 785616 | 231087 |
| 5 | 117676 | 57766 | 144165 | 33171 |
| 6 | 4834 | 0 | 41249 | 24445 |
| 7 | 1807 | 0 | 2053 | 0 |

**Table S4:** Variant calling statistics for ZymoBIOMICS Microbial Community DNA Standard rRNA amplicon libraries. Called variants have been divided into two groups for each data type (ONT UMI, PB CCS, PB UMI): - The variants best matching the references (best variants) and all the other variants (spurious variants). For each group the number of variants and number of errors is listed and the amount of the total data that was used to generate the variants have been calculated.

|  | Best variants |  | Spurious variants |  |
| --- | --- | --- | --- | --- |
| Data type | Number of variants<br>[0 error/1 error] | Fraction of total data<br>[%] | Number of variants<br>[1-5 error/> 5 error] | Fraction of total data<br>[% spurious/<br>% chimeras] |
| ONT UMI | 41/2 | 99.00 | 34/0 | 1/0 |
| PB CCS | 43 | 92.58 | 502/193 | 6.99/0.43 |
| PB UMI | 43 | 99.82 | 2/0 | 0.18/0 |

**Table S5:**
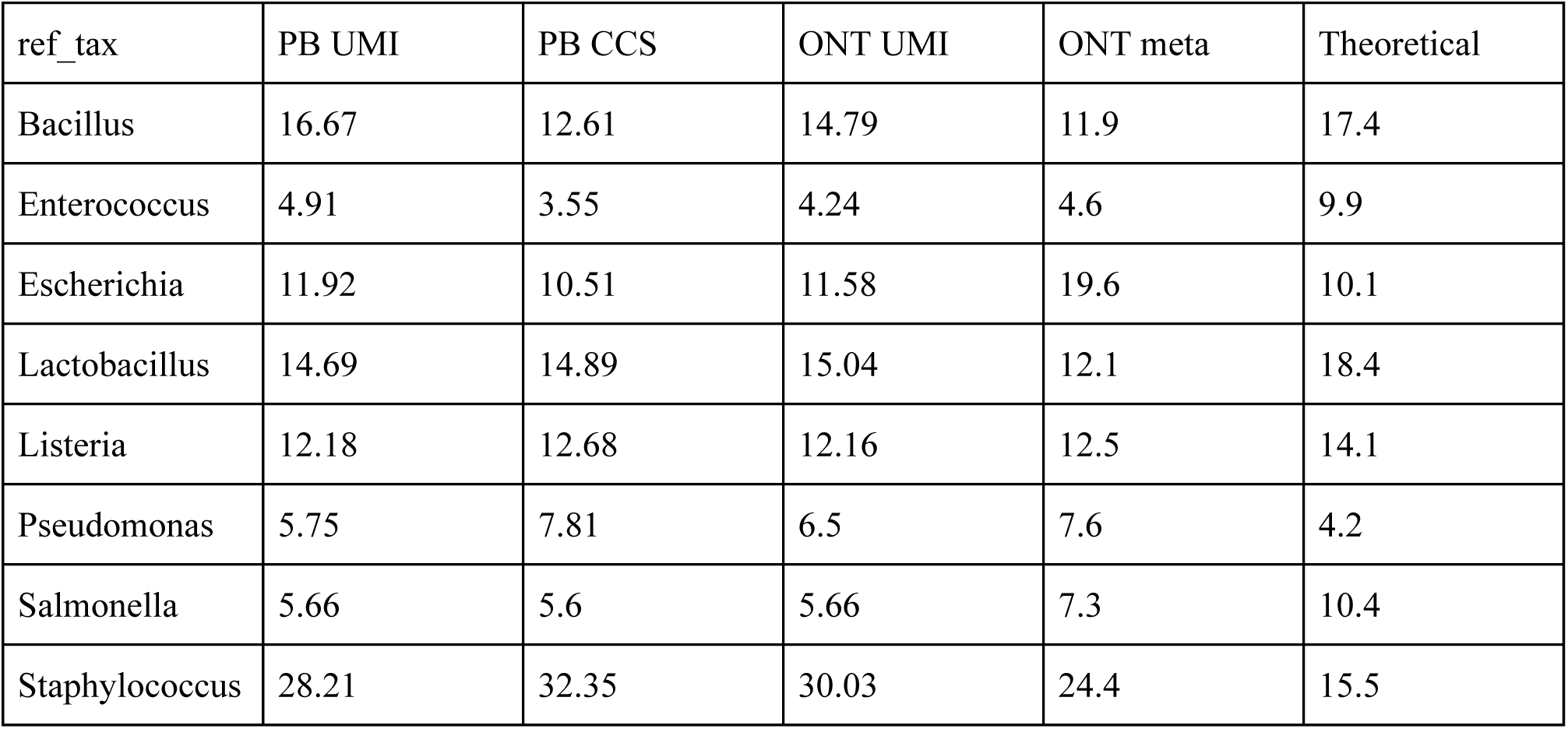
rRNA operon relative abundance estimates. rRNA relative abundance estimated for the different rRNA amplicon data sets (PB UMI, PB CCS, ONT UMI) and compared with abundances estimated from metagenome sequencing data (> 5000 bp used, ONT meta) and the theoretical abundance provided by the vendor. The relative abundance estimates of the mock community was skewed, in the same direction, for all rRNA data types (see Figure S9). If the skew was caused by general PCR bias, we would expect the UMI datasets to be different compared to the CCS dataset, but this is not the case. This indicates the skew originates from the gene specific primers and/or initial template accessibility. Many factors can possibly contribute to the observed skew - most notable are length dependent PCR efficiency (Figure S10), reference dependent DNA fragment size distribution (Figure S11), different growth states (Figure S12), and operon dependent nucleotide composition. The rRNA relative abundance estimated from the ONT metagenome data (ONT meta), is more similar to the rRNA amplicon data, indicating the DNA fragment length plays a major role in the observed relative abundance. A mock community with more defined fragment lengths and genome coverage is needed to evaluate whether the relative abundance estimates of rRNA operons can be applied effectively in microbial ecology.

| ref_tax | PB UMI | PB CCS | ONT UMI | ONT meta | Theoretical |
| --- | --- | --- | --- | --- | --- |
| Bacillus | 16.67 | 12.61 | 14.79 | 11.9 | 17.4 |
| Enterococcus | 4.91 | 3.55 | 4.24 | 4.6 | 9.9 |
| Escherichia | 11.92 | 10.51 | 11.58 | 19.6 | 10.1 |
| Lactobacillus | 14.69 | 14.89 | 15.04 | 12.1 | 18.4 |
| Listeria | 12.18 | 12.68 | 12.16 | 12.5 | 14.1 |
| Pseudomonas | 5.75 | 7.81 | 6.5 | 7.6 | 4.2 |
| Salmonella | 5.66 | 5.6 | 5.66 | 7.3 | 10.4 |
| Staphylococcus | 28.21 | 32.35 | 30.03 | 24.4 | 15.5 |

**Tabel S6:** Cost estimates for different UMI library preparations.

|  | ONT UMI (USD) | PB CCS (USD) | PB UMI (USD) |
| --- | --- | --- | --- |
| Amplicon library | 35 | 15 | 35 |
| Sequencing library | 170 | 485 | 485 |
| Sequencing | 900 | 4032 | 4032 |
| Basecalling/CCS (Cloud) | 65 | 125 | 125 |
| UMI processing (Cloud) | 65 | - | 21 |
| Total | 1235 | 4678 | 4698 |
| See supplementary file '2019-12-17_cost_overview.xlsx' for calculations. |  |  |  |

**Tabel S7:** Time estimates for preparing UMI libraries with the different sequencing platforms.

|  | ONT UMI (hr) | PB CCS (hr) | PB UMI (hr) |
| --- | --- | --- | --- |
| Amplicon library | 6 | 4 | 6 |
| Sequencing library | 2 | 3 | 3 |
| Sequencing | 48 | 35 | 35 |
| Basecalling/CCS (Cloud) | 10 | 24 | 24 |
| UMI processing (Cloud) | 8 | - | 4 |
| Outsourcing | - | (336)<sup>1</sup> | (336)<sup>1</sup> |
| Total time | 74 | 66(402) | 72(408) |
| <sup>1</sup> Outsourcing turnaround time for sequencing according to own experience. |  |  |  |

**Tabel S8A:**
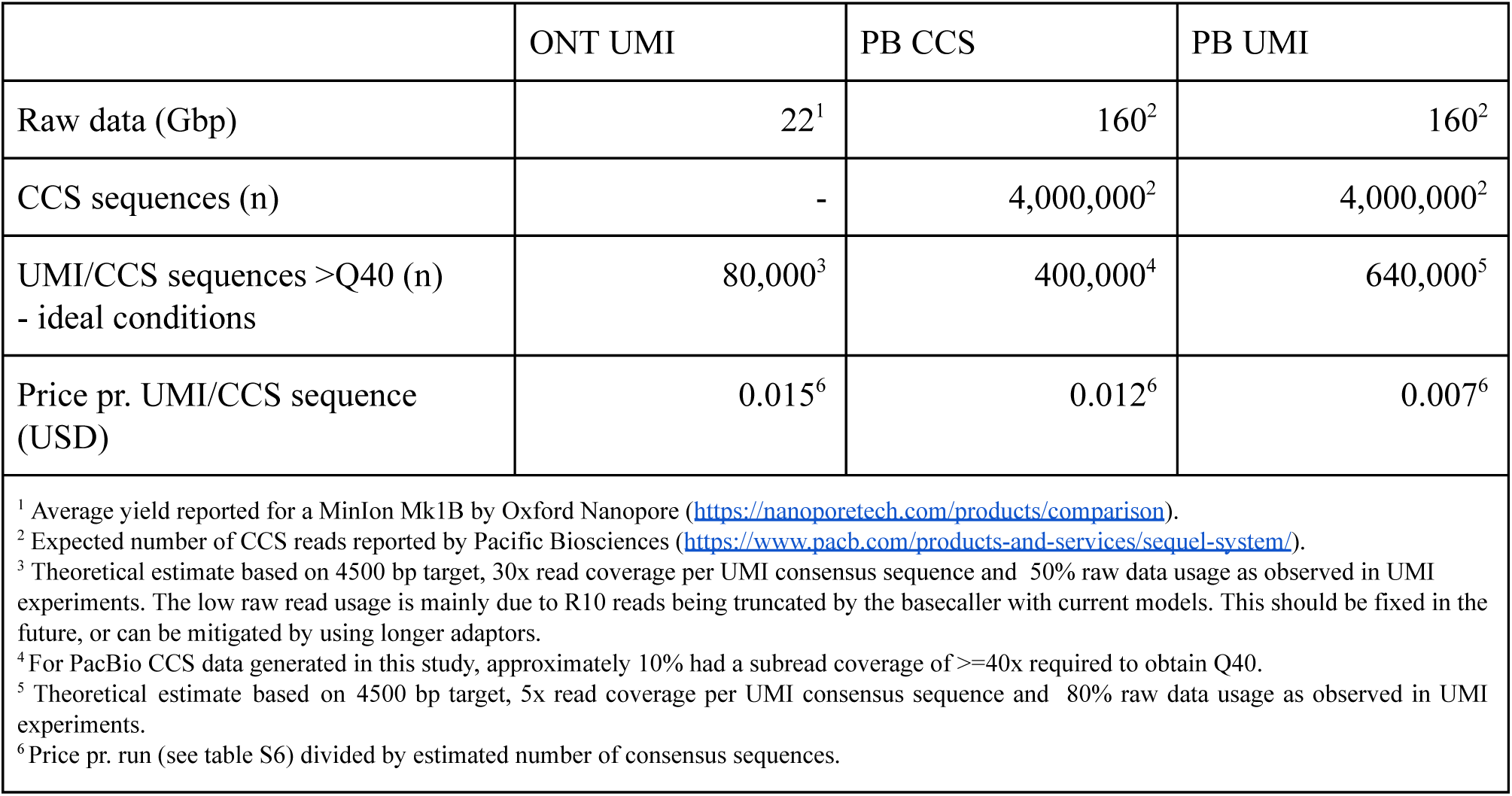
Yield and price estimates for bacterial rRNA operon sequencing (∼4500 bp) with different sequencing strategies under ideal conditions. Ideal conditions assumes yields in line with what is promised by the platform manufactures, and ideal number of template molecules have been used in the PCR. Furthermore, it is assumed that PCR amplification is the same for all molecules.

**Tabel S8B:** Yield and price estimates for bacterial rRNA operon sequencing (∼4500 bp) with different sequencing strategies as observed in this study under non optimal conditions. The number of template molecules was not optimized for yield in this study, and the sequencing runs themselves cannot be viewed as representative. More runs should be performed to calculate meaningful averages. It is therefore not meaningful to compare these yields and prices directly.

|  | ONT UMI | PB CCS | PB UMI |
| --- | --- | --- | --- |
| Raw data (Gbp) | 19 | 161 | 161 |
| CCS sequences (n) | - | 1,908,772 | 1,908,772 |
| UMI/CCS sequences >Q40 (n)<br>- ideal conditions | 23,365 | 135,823 | 39,678 |
| Price pr. UMI/CCS sequence (USD) | 0.053 | 0.034 | 0.118 |

**Table S9:**
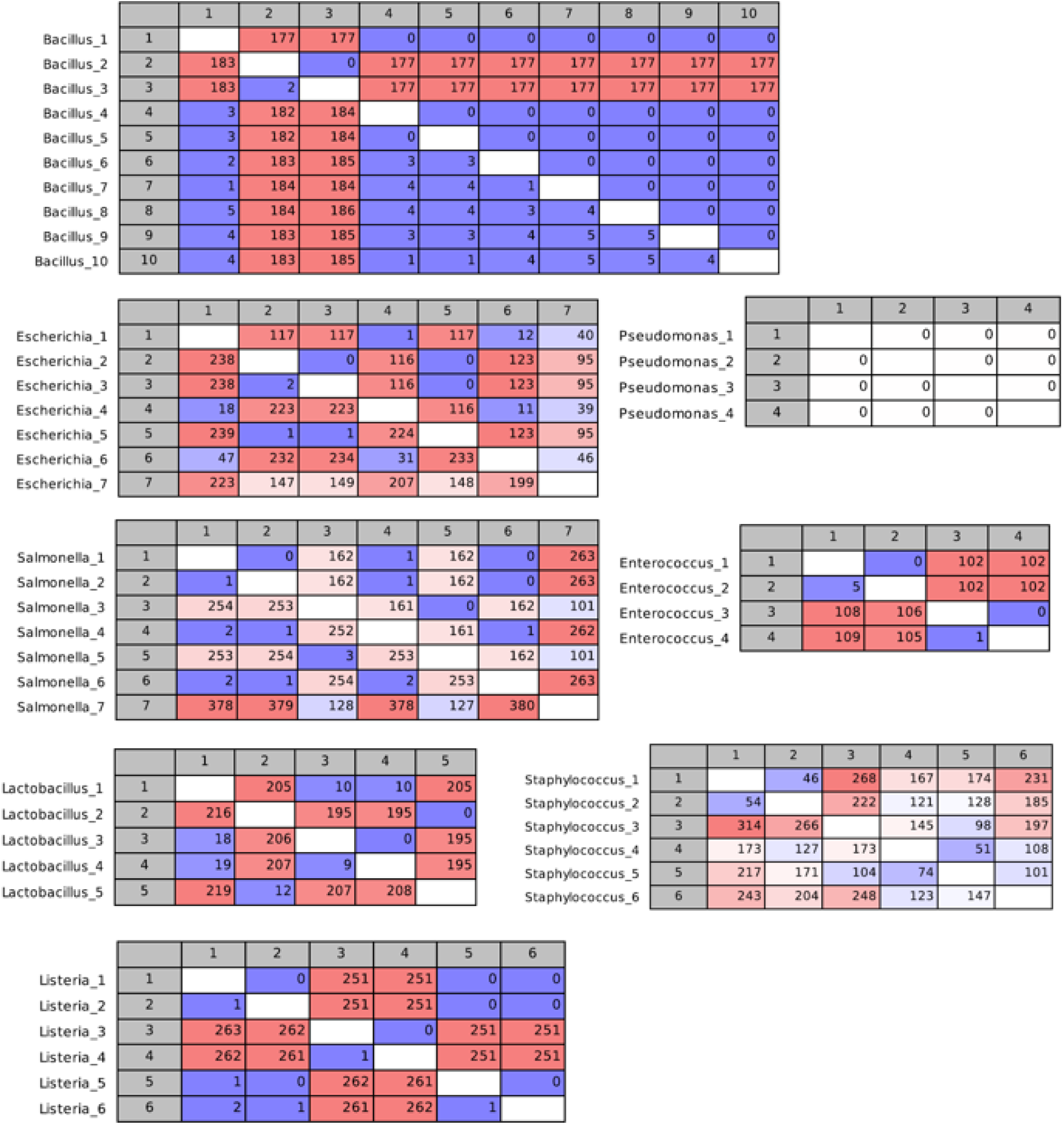
Difference between intra species rRNA operons. Each table show intra species difference between rRNA operons. Below the diagonal is total differences and above is total indels. The analysis was performed on the curated rRNA operons from the ZymoBIOMICS Microbial Community DNA Standard using CLC genomics workbench v9.5.5 (Qiagen) using the ‘Create Alignment’ tool (Gap open cost = 10.0, Gap extension cost = 1.0, End gap cost = Free, Alignment mode = Very accurate (slow), Redo alignments = No, Use fixpoints = No) and the ‘Create pairwise comparison’ tool (default settings).

**Table S10:**
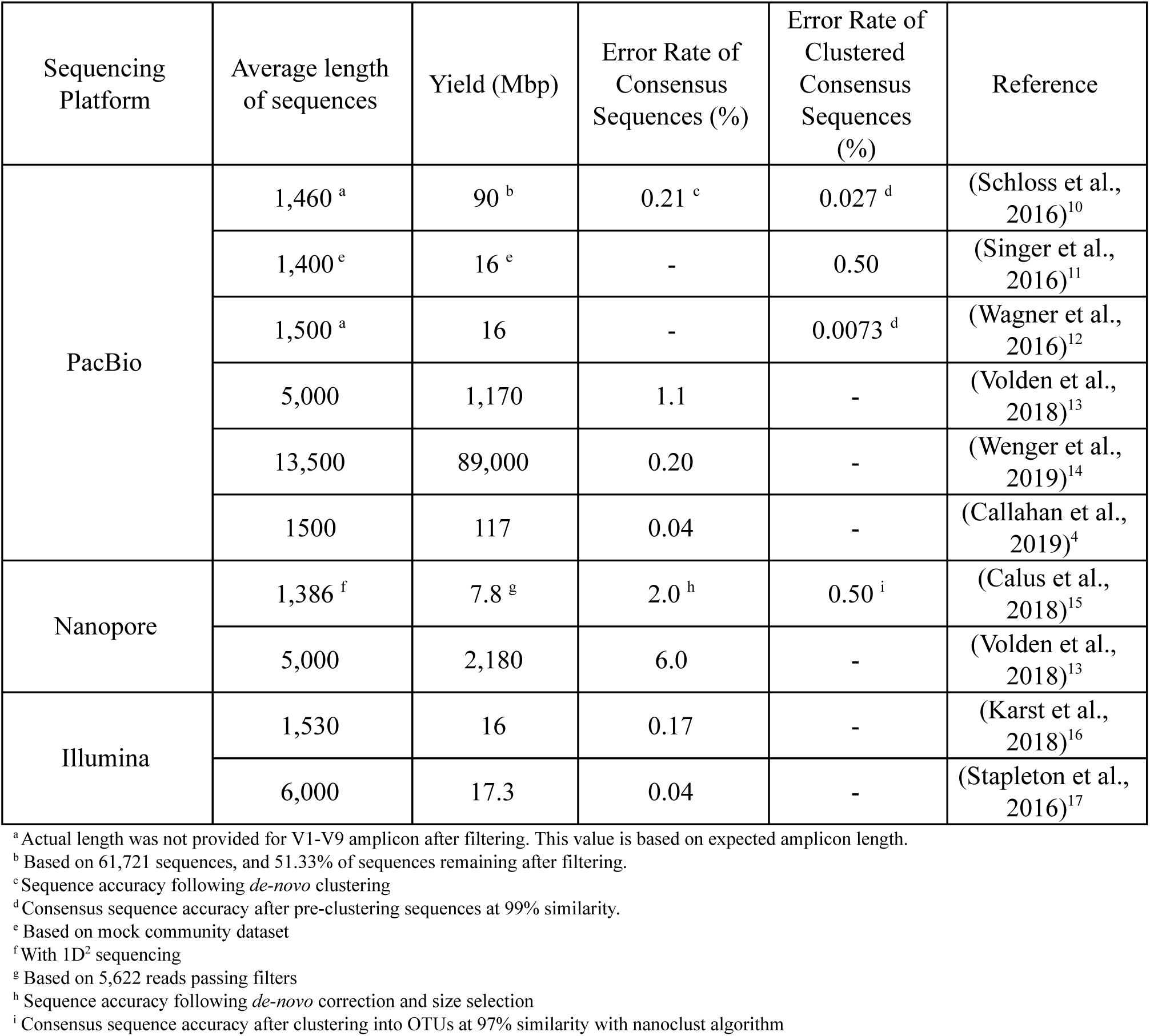
Overview of high-accuracy long amplicon sequencing from the literature.

**Table S11:** Estimation of mismatches between primers and rRNA operon sequences.

| Species | 5' primer hit | 5' primer error (n) | 3' primer hit | 3' primer error (n) |
| --- | --- | --- | --- | --- |
| Bacillus_1 | 27f | 0 | 2490r | 0 |
| Bacillus_10 | 27f | 0 | 2490r | 0 |
| Bacillus_2 | 27f | 0 | 2490r | 0 |
| Bacillus_3 | 27f | 0 | 2490r | 0 |
| Bacillus_4_5 | 27f | 0 | 2490r | 0 |
| Bacillus_6 | 27f | 0 | 2490r | 0 |
| Bacillus_7 | 27f | 0 | 2490r | 0 |
| Bacillus_8 | 27f | 0 | 2490r | 0 |
| Bacillus_9 | 27f | 0 | 2490r | 0 |
| Enterococcus_1 | 27f | 0 | 2490r | 0 |
| Enterococcus_2 | 27f | 0 | 2490r | 0 |
| Enterococcus_3 | 27f | 0 | 2490r | 0 |
| Enterococcus_4 | 27f | 0 | 2490r | 0 |
| Escherichia_1 | 27f | 0 | 2490r | 0 |
| Escherichia_2 | 27f | 0 | 2490r | 0 |
| Escherichia_3 | 27f | 0 | 2490r | 0 |
| Escherichia_4 | 27f | 0 | 2490r | 0 |
| Escherichia_5 | 27f | 0 | 2490r | 0 |
| Escherichia_6 | 27f | 0 | 2490r | 0 |
| Escherichia_7 | 27f | 0 | 2490r | 0 |
| Lactobacillus_1 | 27f | 0 | 2490r | 0 |
| Lactobacillus_2 | 27f | 0 | 2490r | 0 |
| Lactobacillus_3 | 27f | 0 | 2490r | 0 |
| Lactobacillus_4 | 27f | 0 | 2490r | 0 |
| Lactobacillus_5 | 27f | 0 | 2490r | 0 |
| Listeria_1 | 27f | 0 | 2490r | 0 |
| Listeria_2_5 | 27f | 0 | 2490r | 0 |
| Listeria_3 | 27f | 0 | 2490r | 0 |
| Listeria_4 | 27f | 0 | 2490r | 0 |
| Listeria_6 | 27f | 0 | 2490r | 0 |
| Pseudomonas_1_2_3_4 | 27f | 0 | 2490r | 0 |
| Salmonella_1 | 27f | 0 | 2490r | 0 |
| Salmonella_2 | 27f | 0 | 2490r | 0 |
| Salmonella_3 | 27f | 0 | 2490r | 0 |
| Salmonella_4 | 27f | 0 | 2490r | 0 |
| Salmonella_5 | 27f | 0 | 2490r | 0 |
| Salmonella_6 | 27f | 0 | 2490r | 0 |
| Salmonella_7 | 27f | 0 | 2490r | 0 |
| Staphylococcus_1 | 27f | 0 | 2490r | 0 |
| Staphylococcus_2 | 27f | 0 | 2490r | 0 |
| Staphylococcus_3 | 27f | 0 | 2490r | 0 |
| Staphylococcus_4 | 27f | 0 | 2490r | 0 |
| Staphylococcus_5 | 27f | 0 | 2490r | 0 |
| Staphylococcus_6 | 27f | 0 | 2490r | 0 |

**Table S12:** Data overview.

| Experiment | Data type | ENA Accession |
| --- | --- | --- |
| ENA study | - | PRJEB32674 |
| Zymo Mock rRNA operon UMI amplicon data generated with ONT MinION and R9.4.1 flowcell (see <sup>18</sup> for materials and methods) | Raw Nanopore reads (fastq) | ERR3336963 |
|  | Raw Nanopore reads (fast5) | ERR3336964 |
|  | UMI consensus sequences (fasta) | ERZ940787 |
|  | Variants consensus sequences (fasta) | ERZ940796 |
| Zymo Mock rRNA operon UMI amplicon data generated with ONT MinION and R10 flowcell (see materials and methods) | Raw Nanopore reads (fastq) |  |
|  | Raw Nanopore reads (fast5) |  |
|  | UMI consensus sequences (fasta) |  |
|  | Variants consensus sequences (fasta) |  |
| Zymo Mock rRNA operon UMI amplicon data generated with PacBio Sequel II and PacBio Sequel II 8M flowcell in CCS mode (see materials and methods) | Raw subreads (bam) |  |
|  | CCS reads (fastq) |  |
|  | UMI consensus sequences (fasta) |  |
|  | Variants consensus sequences (fasta) |  |
| Escherichia coli str. K-12 substr. MG1655 data generated with ONT MinION and R10 flowcell (see materials and methods) | Raw Nanopore reads (fastq) |  |
|  | Raw Nanopore reads (fast5) |  |
|  | UMI consensus sequences (fasta) |  |
|  | - |  |
| American Gut Project<sup>19</sup> fecal rRNA operon data generated with PacBio Sequel II and PacBio Sequel II 8M flowcell in CCS mode (see materials and methods) | CCS reads (fastq) with sample identifiers in the header |  |

**Table S13:** Sequencing yield statistics

|  | ONT UMI | PB CCS | PB UMI |
| --- | --- | --- | --- |
| Raw reads (n) | 4,412,447 | 36,630,361 | 1,908,772 |
| Raw bases (Mbp) | 18,888 | 161,430 | 8,485 |
| UMI binned reads (n) | 1,131,157 | - | 1,576,585 |
| UMI binned bases (Mbp) | 4,957 | - | 6,900 |
| Consensus reads (n) | 38,926 | 1,908,772 | 39,678 |
| Consensus bases (Mbp) | 170 | 8,485 | 173 |
| >Q40 Consensus reads (n) | 23,365 | 135,823 | 39,678 |
| >Q40 Consensus bases (Mbp) | 102 | 459 | 173 |

'Raw' is sequencing data directly after basecalling before any filtering. 'UMI binned' is data that has been quality filtered and been successfully assigned to a specific UMI bin. 'Consensus' is data in consensus sequence form, either number of CCS consensus reads or number UMI consensus reads. '>Q40 Consensus' is data in consensus sequence form filtered based on coverage to obtain >Q40 data.

